# Negative Affect Induces Rapid Learning of Counterfactual Representations: A Model-based Facial Expression Analysis Approach

**DOI:** 10.1101/560011

**Authors:** Nathaniel Haines, Olga Rass, Yong-Wook Shin, Joshua W. Brown, Woo-Young Ahn

**Affiliations:** The Ohio State University; Indiana University Bloomington; University of Ulsan; Seoul National University

**Keywords:** counterfactual thinking, regret, disappointment, reinforcement learning, Bayesian analysis

## Abstract

Whether we are making life-or-death decisions or thinking about the best way to phrase an email, counterfactual emotions including regret and disappointment play an ever-present role in how we make decisions. Functional theories of counterfactual thinking suggest that the experience and future expectation of counterfactual emotions should promote goal-oriented behavioral change. Although many studies find empirical support for such functional theories, the generative cognitive mechanisms through which counterfactual thinking facilitates changes in behavior are underexplored. Here, we develop generative models of risky decision-making that extend regret and disappointment theory to experience-based tasks, which we use to examine how people incorporate counterfactual information into their decisions across time. Further, we use computer-vision to detect positive and negative affect (valence) intensity from participants’ faces in response to feedback, which we use to explore how experienced emotion may correspond to cognitive mechanisms of learning, outcome valuation, or exploration/exploitation—any of which could result in functional changes in behavior. Using hierarchical Bayesian modeling and Bayesian model comparison methods, we found that a model assuming: (1) people learn to explicitly represent and subjectively weight counterfactual outcomes with increasing experience, and (2) people update their counterfactual expectations more rapidly as they experience increasingly intense negative affect best characterized empirical data. Our findings support functional accounts of regret and disappointment and demonstrate the potential for generative modeling and model-based facial expression analysis to enhance our understanding of cognition-emotion interactions.

## Theories of counterfactual thinking: Core concepts and proposed functions

Counterfactual thinking is the act of mentally simulating “what could have been”, had either chance produced a different state of the world, or had one made a different decision altogether. From high-stakes decisions such as choosing between life-or-death medical procedures, to every-day questions like choosing what to eat for lunch, people often make decisions based on the probable (and often subjective) consequences of their actions. Functional theories of counterfactual thinking suggest that mental simulation and comparison of these consequences elicits emotional reactions, which then serve to motivate goal-oriented changes in behavior.

*Regret* is one such emotion, which is characterized by an aversive subjective state that occurs when we make one choice and then later wish to have made an alternative choice (Kahneman & Miller, 1986). In fact, some studies show that regret is one of the most frequently experienced and discussed emotions in everyday situations (Shimanoff, 1984), and people who experience regret either most often and/or most intensely endorse more severe symptoms of anxiety and depression in addition to lower life satisfaction than their peers (Kocovski, Endler, Rector, & Flett, 2005; Lecci, Okun, Karoly, 1994, 1994; Monroe, Skowronski, Macdonald, & Wood, 2005). Despite being associated with these negative long-term outcomes, many people look back on regretted decisions with appreciation, which is not true of other negative emotions like anger, jealousy, anxiety, guilt, and boredom (Saffrey, Summerville, & Roese, 2008). These seemingly paradoxical findings are reconciled by functional theories of counterfactual thinking including decision justification theory and regret regulation theory (Connolly & Zeelenberg, 2002; Zeelenberg & Pieters, 2007), wherein regret is hypothesized to motivate goal-oriented behavioral change by signaling us to avoid making choices in the presence of more optimal alternatives (see also Epstude & Roese, 2008; Roese, 1994). Indeed, the intensity of regret that we experience is proportional to how active our role is in the regret-inducing decision, how justifiable our decision is, and the quality of our decision process (e.g., Inman & Zeelenberg, 2002; Pieters & Zeelenberg, 2005). Further, we experience the most regret in domains where we have opportunities for corrective action (Roese & Summerville, 2005). However, not all forms of negatively-valenced counterfactual thinking lead to long-term behavioral changes, suggesting that regret serves a specific function in how we decide among multiple risky choices (e.g., Zeelenberg et al., 1998).

*Disappointment* is another example, which occurs when comparing an obtained outcome to what could have been obtained given an alternative state of the world (Kahneman & Miller, 1986). Like regret, disappointment is experienced as aversive, yet it is evaluated favorably in retrospect (Saffrey, Summerville, & Roese, 2008). Individual differences in how children regulate experienced disappointment predicts behavioral problems including symptoms of attention-deficit/hyperactivity disorder and oppositional/aggressive behavior (Cole, Waxler, & Smith, 1994), and adults who experience more frequent disappointing life events tend to show higher rates of depression (Goodyer, Herbert, Tamplin, Secher, & Pearson, 1997). Unlike regret, disappointment appears to play a more primary role in modifying how people evaluate situational outcomes or one-off events as opposed to promoting specific changes in behavior (e.g., Markman, Gavanski, Sherman, & McMullen, 1993; Zeelenberg et al., 1998). For example, experimental evidence suggests that disappointment can lead one to feel worse about outcomes they receive when there are salient, better outcomes—and alternatively, people feel better about outcomes they receive when they could have received worse outcomes (e.g., Ordóñez, Connolly, & Coughlan, 2000). Therefore, the functional value of disappointment appears to be related to coping and reacting to momentary outcomes that indirectly facilitate more long-term positive gains, whereas regret is more directly related to long-term changes in behavior.

Although the empirical studies described above consistently implicate counterfactual emotions in goal-oriented behavioral change, there is a relative absence of formal generative models guiding research on how people learn to represent and therefore modify their behavior based on counterfactual information. A deeper understanding of such mechanisms may help us determine why regret can sometimes lead to positive behavioral changes (e.g., corrective action), whereas other times it can lead to negative behavioral outcomes (e.g., depression, anxiety, lower life satisfaction, etc.). Similarly, identifying such mechanisms may elucidate the functional role of disappointment, thus providing an explanation for prospective links between disappointing events and the behavioral problems described above. Below, we review existing generative models of counterfactual thinking in addition to shortcomings that limit their use for understanding functional theories of counterfactual thinking.

## Theories of counterfactual thinking: Formal generative models

Generative models of counterfactual thinking date back to early work on regret and disappointment theory (Bell, 1982; 1985; Loomes & Sugden, 1982b; 1986; see also Savage, 1951). According to regret theory, counterfactual comparisons between outcomes *across* actions lead to cognitive-emotional states of *regret* or *rejoicing* if the comparison is negative or positive in value, respectively. With only two possible actions, expected regret/rejoice for a given action *i* ∈ {1,2} is formulated as the sum of all possible differences between each of its possible outcomes *j* versus each possible counterfactual outcome *k* of the foregone action *i*′, where each difference is weighted by its joint probability of occurring:

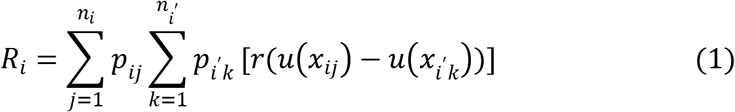

Here, *n_i_* and *n*_*i*′_ represent the total number of outcomes for action *i* and foregone action *i*′, respectively, *u*(*x*) is a traditional utility function that assigns subjective value to outcomes independently, and *r*(*x*) is a utility function that acts on the counterfactual difference, which can differ between regret and rejoice outcomes (i.e. when the difference is positive and negative, respectively). Panels 1 and 2 of Figure 1 demonstrate the logic of Equation 1.

**Figure 1.**
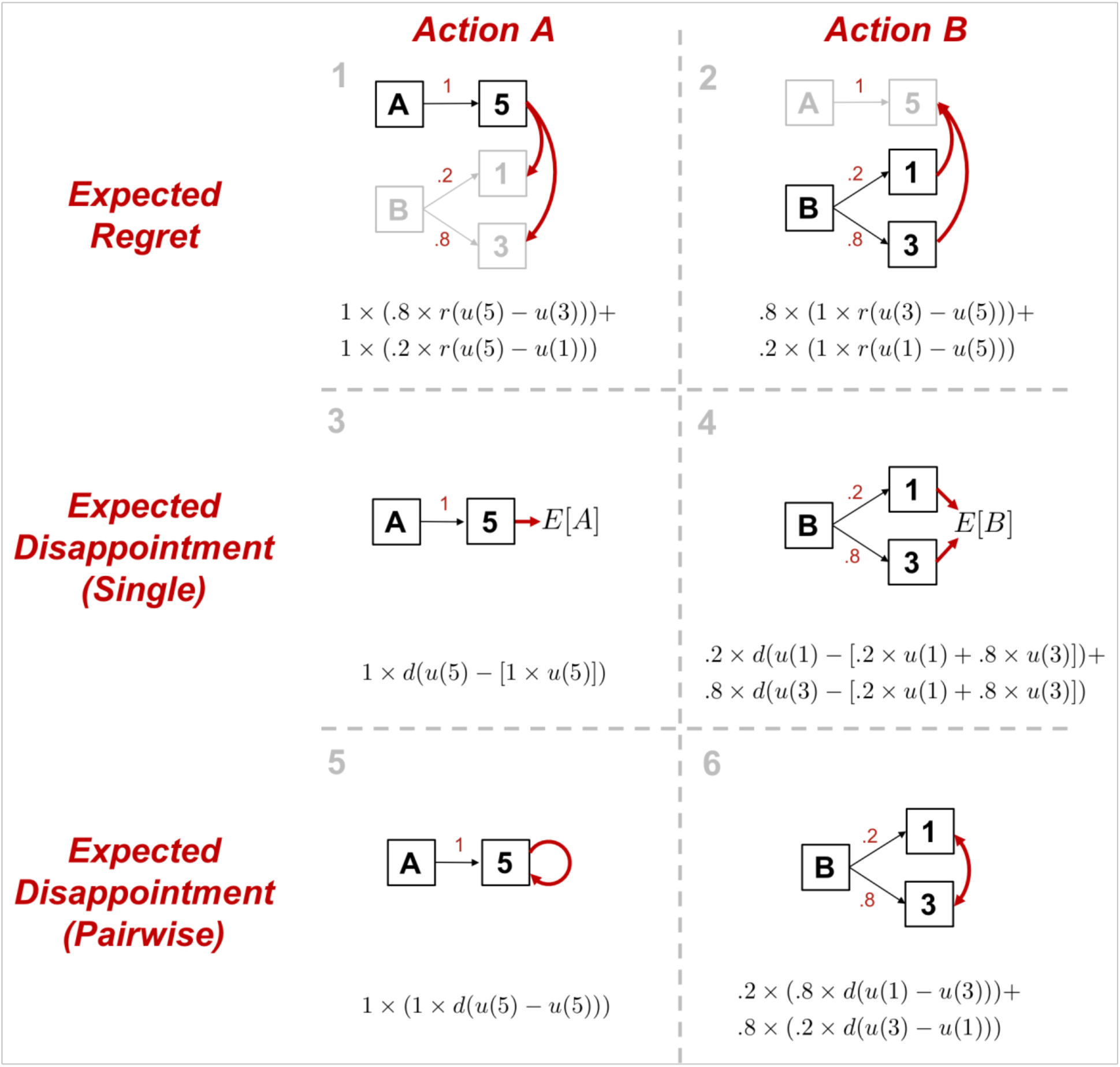
Generative models of counterfactual thinking Here we represent generative models counterfactual thinking, including those derived from regret and disappointment theory. The choice represented above is the following: “*Do you prefer action A: $5 with Pr(1); or action B: $1 with Pr(.2), $3 with Pr(.8)?*”. Panels 1-2, 3-4, and 5-6, represent Equations 1, 2, and 3 in the main text, respectively. Note that for panels 3 and 5, we represent expected disappointment as a comparison of an outcome with its itself, given it is the only possibility. This is equivalent to assuming that expected disappointment is 0.

Although Equation 1 may at first appear quite complex, the intuition behind it is simple— that people generate a regret expectation by thinking of what it would feel like to receive each potential consequence of an action, relative to each consequence of a salient alternative action. Here, salience is determined by the objective probability of the consequence occurring. This explicit representation of relative differences in the utility of outcomes across actions can capture shortcomings of competing theories (e.g., expected utility theory) while maintaining a simple form with relatively few assumptions compared to other models (see Loomes & Sugden, 1982a; 1982b). An example is the correlation effect, which manifests as a change in expected-value maximization behavior when the outcomes across actions are correlated (Diederich & Busemeyer, 1999; Erev et al., 2017; Grosskopf, Erev, & Yechiam, 2006). More importantly for the current study, the formal specification of regret theory makes specific commitments regarding how people use information about the consequences of their actions to guide future behavior, which can be tested against empirical data.

Disappointment theory was later developed and incorporated into models of regret (Bell, 1985; Loomes & Sugden, 1986). According to disappointment theory, counterfactual comparisons between outcomes *within* actions lead to states of *disappointment* or *elation* if the comparison is negative or positive in value, respectively. Historically, expected disappointment/elation has been formulated in various different ways. Both Bell (1985) and Loomes and Sudgen (1986) assumed that the expected disappointment/elation for a given action *i* is the weighted sum of the difference between each possible outcome resulting from that action versus its expected subjective utility (*E*[*u*(*X_i_*)]):

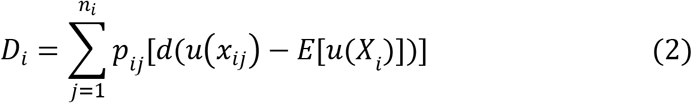

Here, *d*(*x*) is a utility function that acts on the difference between outcome *x_ij_* and the action *i*’s expectation, which can differ between disappointment and elation comparisons (i.e. when the difference is positive and negative, respectively). Experienced disappointment and elation can therefore be thought of as prediction errors, where expected disappointment and elation are then weighted sums of all possible prediction errors. Note that *D_i_* = 0 so long as the disappointment utility function is symmetric across positive and negative prediction errors (i.e. *d*(*u*(*x_ij_*) − *E*[*u*(*X_i_*)]) = *d*(*E*[*u*(*X_i_*)] − *u*(*x_ij_*))). Panels 2 and 3 of Figure 1 demonstrate the logic of Equation 2.

Equation 2 makes predictions regarding how experienced outcomes should relate to experienced positive/negative affect (or valence)—specifically, valence is assumed to result from differences between experienced and expected choice outcomes, such that even strongly valenced outcomes (e.g., losing a large sum of money, getting a big promotion) are not predicted to induce strong emotional states if more extreme outcomes were expected (e.g., a $5,000 salary raise may be valued negatively if a $10,000 raise was expected). However, as shown convincingly by Ordóñez et al. (2000), people tend to compare outcomes in a pairwise manner, and not to a single overall expectation per se. To better capture this tendency, Delquié and Cillo (2006) reformulated disappointment theory such that expected disappointment/elation for a given action *i* is the expectation over all pairwise differences between outcomes within an action *i*:

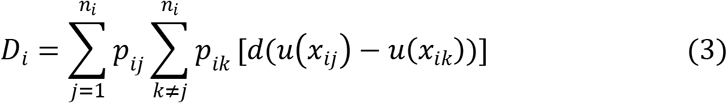

Intuitively, Equation 3 assumes that people generate disappointment expectations by thinking of what it would be like to receive each possible outcome relative to all other possible outcomes when engaging in a specific action, where each pairwise comparison is weighted by its joint probability of occurring. As in the expected value formulation, the pairwise formulation of disappointment theory allows for *d*(*x*) to vary depending on the sign of the difference, such that *d*(*u*(*x_ij_*) − *u*(*x_ik_*)) ≠ *d*(*u*(*x_ik_*) − *u*(*x_ij_*)). Somewhat unintuitively, common risky decision-making models including the risk-value, rand dependent utility, and mean-variance models can be derived directly from the pairwise formulation of disappointment theory, thus allowing it to provide a psychologically plausible explanation for how people incorporate risk and uncertainty into their decisions (see Delquié & Cillo, 2006). Panels 5 and 6 of Figure one demonstrate the logic of Equation 3.

Comparing Equation 3 to Equation 1, we can see that the pairwise disappointment formulation is identical in form to classic regret theory—the only difference being that regret versus disappointment involve comparisons of outcomes *across* versus *within* actions, respectively (in addition to the regret and disappointment specific utility functions). The pairwise difference formulations of regret and disappointment theory therefore provide a consistent, unified account of counterfactual thinking. Indeed, models based on the pairwise difference formulations of regret and disappointment theory have been used to explain behavior in a variety of different contexts—both neuroimaging studies and those collecting subjective reports of emotion during regret- and disappointment-inducing tasks support the notion that emotional states of regret and disappointment arise from pairwise comparisons between outcomes (e.g., Coricelli et al., 2005; Lohrenz, McCabe, Camerer, & Montague, 2007; Mellers et al., 1997; 1999; Ordóñez et al., 2000). The empirical success of regret and disappointment theory in both explaining choice behavior and subjective reports of emotion makes them ideal for exploring functional theories of counterfactual thinking.

## Using regret and disappointment theory to explain changes in behavior

Despite the success of regret and disappointment theory in explaining both choice behavior and subjective emotional states, there is one key shortcoming that precludes their use for understanding and testing functional theories of counterfactual thinking. Specifically, most previous studies have used description-based tasks and static models, wherein the outcomes and associated probabilities of each action are known a priori and assumed to be constant across trials (Coricelli et al., 2005; Mellers et al., 1997; 1999; but see Ahn et al., 2012). A consequence of this design choice is that behavior itself is assumed to be constant across time, which is in direct opposition to the supposed functional role of counterfactual thinking in changing future behavior. Further, it is inconsistent with empirical evidence showing that people do, in fact, change their behavior when experiencing regretful outcomes in description-based tasks (e.g., “*would you rather have $3 with Pr(1), or $4 with Pr(.8), 0 otherwise?*”), so long as they receive feedback on their choices (e.g., Erev et al., 2017). In studies that do require people to learn the outcomes and associated probabilities of their actions from a state of no knowledge (e.g., “bandit” learning tasks), proposed models often deviate quite substantially from Equations 1-3, instead using principles derived from either reinforcement- or instance-based learning theories. As implemented, such models learn to represent a single running expectation of expected value, regret, or disappointment for each outcome, whereas regret and disappointment theory assume an explicit representation of and pairwise comparisons between each outcome (e.g., Erev & Roth, 2014; Erev et al., 2014; 2017; Lohrenz, McCabe, Camerer, & Montague, 2007; Yechiam & Rakow, 2012).

A related concern is the so-called description-experience gap—people act *as if* they overweight low probabilities when making description-based choices yet underweight low probabilities when making experience-based decisions (e.g., Barron & Erev, 2003; Griffiths & Tenenbaum, 2006; Hertwig & Erev, 2009; Hertwig, Barron, Weber, & Erev, 2004; Ungemach, Chater, & Stewart, 2009; Wulff, Mergenthaler-Canseco, & Hertwig, 2018). Therefore, despite regret and disappointment showing robust effects across a number of different description- and experience-based decision paradigms (e.g., Avrahami, Kareev, & Hart, 2014; Hart, Avrahami, Kareev, Todd, 2015; Kareev, Avrahami, & Fielder, 2014; Marchiori & Warglein, 2008; Rakow, Newell, Wright, 2015), the extent to which the principles underlying regret and disappointment theory can generalize to pure experience-based paradigms is less clear.

In summary, many models of counterfactual thinking being proposed throughout the psychological, cognitive, and brain sciences, yet there is a lack of work focused on developing models that retain key assumptions of regret and disappointment theory in a way that makes them amenable for exploring functional theories of counterfactual thinking. The key problem is straightforward—the pairwise difference formulations of regret and disappointment theory have been successful in explaining both neural and subjective responses of experienced emotion, yet they do not include a learning or memory mechanism and therefore cannot explain changes in behavior over time. Conversely, reinforcement- and instance-based learning models of counterfactual thinking can explain changes in behavior, yet they often deviate non-trivially from regret and disappointment theory, and they have been developed without making explicit connections between the cognitive models and experienced emotion. This gap in the literature is a potentially important oversight, given the sophistication of regret and disappointment theory, the supposed functional role of experienced emotion in motivating behavioral change, and the recent movements toward linking traditionally separated components of cognition and emotion (Barrett, 2009; Duncan & Barrett, 2007; Eldar et al., 2016; Etkin et al., 2015; Pessoa, 2008; Pessoa & Adolphs, 2010).

## The current study

Our goals for the current study are two-fold. First, we aim to extend the pairwise formulations of regret and disappointment to contexts wherein peoples’ behavior is expected to change over time as they experience the consequences of their actions (e.g., Ahn et al., 2012). Second, we aim to use real-time measures of emotion to inform model development, which will allow us to better understand how changes in experienced emotion relate to changes in behavior.

To address our first goal, we developed 5 different models, each making different assumptions about how people represent and make decisions based on counterfactual information. Each of these models embodies key principles from previously suggested models in the literature (e.g., explicit representation of outcomes and probabilities, versus singular representations of expected regret). We test these models in two separate studies. In the first study, we use data collected from a large number of participants undergoing various repeated description-based decisions with full-information feedback. In the second study, we apply the same 5 models to data collected in the context of a pure experience-based paradigm with full-information feedback.

To address our second goal, we used computer vision to record participants’ facial expression valence intensity in response to outcome feedback within the experience-based task in study 2. We use trial-level changes in facial expression valence as direct input into the best-fitting cognitive model from studies 1 and 2 to determine if participants’ facial affect in response to feedback relates to changes in either: (1) how quickly they update their expectations, (2) how they assign value to outcomes, or (3) their tendency to explore versus exploit actions. This model-based approach allowed for us to identify relationships between measures of emotion and subsequent changes in behavior in an effort to deepen our understanding of functional theories of counterfactual thinking.

## Study 1 Method

### Participants

Our first study included a combined total of 686 participants’ data collected across both the 2015 and 2018 Choice Prediction Competitions (Erev, Ert, Plonsky, Cohen, & Cohen, 2017; Plonsky, Erev, & Ert, 2017). We excluded trials where participants made choices that did not fit our task inclusion criteria (more detail below in *2.2 Behavioral Task*), but all participants were included.

### Behavioral task

In the Choice Prediction Competition data, all participants completed one of seven sets of 30 gambling games (for a total pool of 210 games) in randomized order. For the current study, we excluded games within each set containing more than two options or payoffs to restrict our focus to 2-choice paradigms. Additionally, we excluded games involving choice ambiguity, where payoffs/probabilities were only partially described. Altogether, these exclusions restricted the number of possible games to 105 (mean [SD] number of games played across participants = 14.9 [5.1]; range = 5-24) (see Supplementary Table 1 for a breakdown of included games).

Each game consisted of 25 trials, where the true payoffs and associated payoff probabilities were shown for each option before a choice was made on each trial (termed repeated description-based choices with full feedback). Each set included a mix of games using: (a) safe vs. risky options and risky versus risky options, and (b) positive and negative outcomes. Additionally, for the first 5 trials within each game, no feedback on choice outcomes was revealed; starting from the 6^th^ trial, “full-information” feedback on both the chosen and forgone outcome was given (see Figure 2 for an example trial). Participants were instructed to make preference-based choices, and that they would be compensated using the real outcome of one randomly selected trial across trials and games.

**Figure 2.**
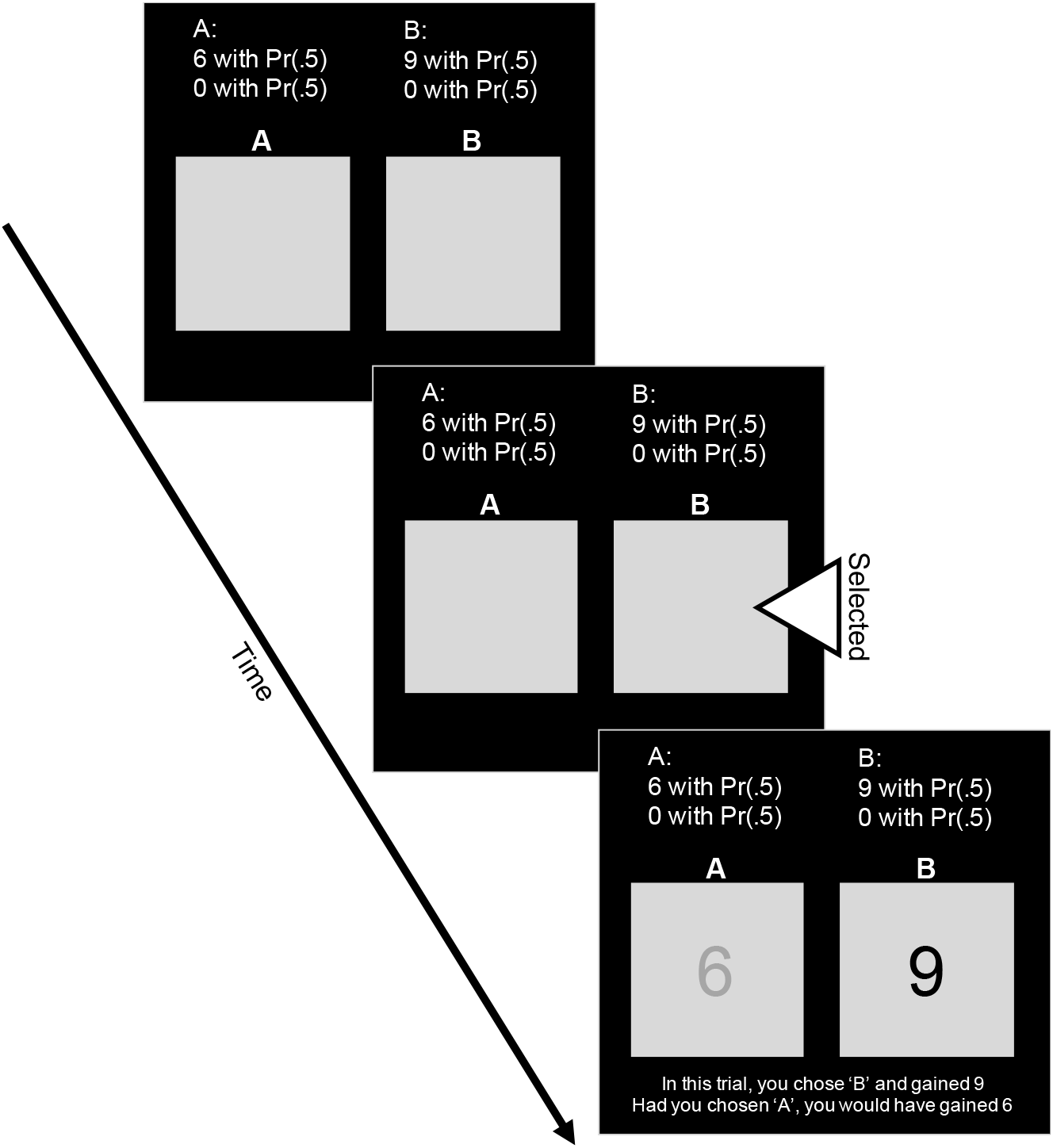
Description-based behavioral task with full-information feedback. In the description-based task, participants were presented with complete information on the outcome distribution of each choice, which remained constant within each game. For the first 5 trials, feedback (i.e. the results of the choice shown in the last panel) was not presented. Starting from the 6^th^ trial, participants received “full information” feedback on the outcome of the chosen and foregone choice outcomes.

### Description-based computational models

We fit a total of 5 different models designed to test different counterfactual learning mechanisms that participants may use to develop a representation of disappointment/regret over time. Each model assumes that participants make choices to maximize their subjective expected pleasure, which varies over time depending on the outcomes they observe with experience. The models are all equivalent in the context of pure description-based decisions with no feedback, and they also all share the same three free parameters: (1) a learning rate, (2) a utility shape parameter, and (3) a choice sensitivity (or inverse temperature) parameter. Therefore, the models only differ in how they assume that a representation of subjective expected pleasure evolves with experience over time. We describe each of the models in detail below, along with a theoretical rationale for why we included them.

#### Counterfactual Representation Learning Model

Of the models we developed, the Counterfactual Representation Learning model most closely resembles traditional decision-theoretic models of disappointment and regret (Bell, 1982; 1985; Loomes & Sugden, 1982; 1986; Mellers et al. 1997; 1999). We assume that people make choices to maximize their subjective expected pleasure (*SEP*). The *SEP* for each action *i* on trial *t* is given by:

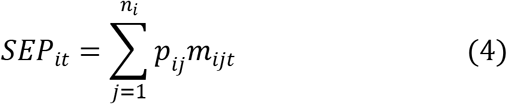

where *p_ij_* indicates the objective, described probability of outcome *j* for action *i*, *n_i_* is the number of possible outcomes for choice option *i*, and *m_ijt_* is the *modified utility*. The modified utility incorporates counterfactual information into the traditional utility function:

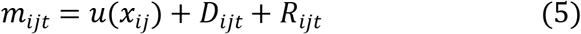

Here, *x_ij_* is the outcome corresponding to *p_ij_* in Equation 4, and *u*(*x*) = *sign*(*x*) ⋅ |*x*|^*w*^ is a utility function that assigns subjective value to outcomes, where *w* (0 < *w* < 1.5) is a person-specific utility shape parameter.

Then, *D_ijt_* is an *expected disappointment* term (or *elation*, depending on if it is negative or positive in value, respectively), which is a weighted sum of the difference between outcome *j* for choice option *i* and each possible alternative outcome *within* option *i*:

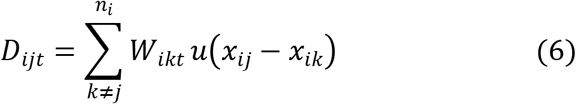

Here, *W_ikt_* is a dynamic “experience weight” for the counterfactual outcome (*x_ik_*), and *u*(*x*) = *sign*(*x*) ⋅ |*x*|*w* is the same utility function from Equation 5. Note that Equation 6 deviates from classic disappointment theory by assuming that the counterfactual terms (*x_ik_*) are weighted by their experience weights (*W_ikt_*), as opposed to their objective probabilities (*p_ik_*), which will allow the model to capture changes in behavior over time (more details below). Further, we follow the pairwise difference formulation of disappointment described in the introduction (see Delquié & Cillo, 2006). Additionally, because we were primarily interested in changes in behavior over time, we simplified the utility functions in Equation 6 relative to the disappointment theory models described in the introduction (see Equation 3). Specifically, we assume a single utility function, controlled by *w*, which is used to assign subjective utility to both individual outcomes and counterfactual differences. Further, we assume that *w* is identical across positive and negative comparisons (i.e. elation and disappointment, respectively).

Due to the simplified utility functions, if the counterfactual experience weight is equivalent to its objective probability (i.e. *W_ikt_* = *p_ik_*), then *D_ijt_* = 0 and expected disappointment drops out of the modified utility term. Similarly, because we initialize all experience weights at 0 (details described below), there is no expected disappointment/elation before participants experience feedback on the outcomes of their choices. Therefore, Equation 6 produces a dynamic disappointment expectation that varies with experience until all experience weights converge to their true values. This model produces behavior similar to a win-stay-lose-shift heuristic (e.g., Worthy, Hawthorne, & Otto, 2013), where actions that have more recently returned their best outcome (i.e. a “win”) are increasingly preferred and vice-versa. Note that it is straightforward to extend the model to include different valuation mechanisms for disappointment versus elation (e.g, a different *ω* parameter for when the difference in Equation 6 is negative versus positive, respectively), which can lead to a non-zero disappointment expectation even when all *W_ikt_* = *p_ik_* (see Delquié & Cillo, 2006).

Finally, *R_ijt_* is an *expected regret* term (or *rejoice*, depending on if it is negative or positive in value, respectively), which is a weighted sum of the difference between each possible outcome *j* of option *i* and each possible outcome *k* of the alternative option *i*′:

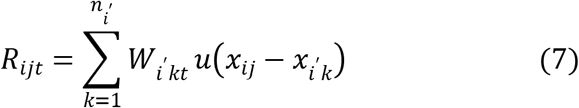

Here, *u*(*x*) is the same utility function as described above, sharing the same shape parameter *w*. Therefore, regret is similar to disappointment, but the counterfactual expectation is computed *across* options as opposed to *within* options as described in the introduction. In Equation 7, we again deviate from classic regret theory by assuming that the regret weight is a dynamic experience weight (*W_ikt_*) as opposed to the objective probability of the counterfactual outcome (cf. Equation 7 above to Equation 1, or to Equations 2-3 in Loomes & Sugden, 1982). Similar to disappointment, the trial-dependent experience weight allows for regret to vary with experience over time, which we describe in more detail below.

Lastly, we use a logistic (i.e. softmax) choice rule to transform subjective expected pleasure (*SEP_it_*) into choice probabilities for each action *i*:

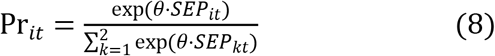

where *θ* is a person-specific and trial-independent choice sensitivity (i.e. inverse temperature) parameter determined by *θ* = 3^*c*^ − 1 that captures how deterministically versus randomly participants make choices according to the differences in *SEP* across options (Yechiam & Ert, 2007). Here, the free parameter is *c* (0 < *c* < 5). In Equation 8, we only sum over two options given that all gambles in our data involve only two options.

The crucial component of the model is the set of counterfactual experience weights (***W***), which allow for behavior to change as a function of experience. For example, when ***W*** = 0 before any feedback is received, both the regret and disappointment terms return 0, and the modified utility reduces to a basic expected utility model similar to prospect theory (albeit without loss aversion or the probability weighting parameters in the current implementation; Kahneman & Tversky, 1979). Conversely, if ***W*** are set to their corresponding objective outcome probabilities (i.e. all *W_ikt_* = *p_ik_*), then the model is akin to traditional regret theory (disappointment drops out, as described above), which is consistent with empirical evidence that regret dominates behavior with increasing experience (e.g., Kareev et al., 2014). We capture learning from experience by assuming that ***W*** = 0 before participants receive experience-based feedback, where ***W*** → ***p*** as participants gain experience. We use the following delta learning rule (a simplified version of the Rescorla-Wagner updating rule; Rescorla & Wagner, 1972) to formalize this learning process:

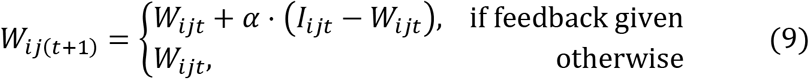

Here, *α* is a learning rate determining how rapidly participants update the experience weight after observing feedback on their choice, and *I_ijt_* is an indicator variable that returns 1 if an outcome was observed during feedback and 0 otherwise. We chose to weight regret by its learned expectedness both to retain consistency with regret and disappointment theory, and because observational and survey-based evidence consistently shows that people experience more regret when they make low-quality or unjustified decisions (e.g., Inman & Zeelenberg, 2002; Pieters & Zeelenberg, 2005). Intuitively, Equation 9 captures this effect by making the regret term more extreme as the regretful outcomes become more subjectively probable. Additionally, our previous work shows that learning rules that explicitly update outcome probabilities can capture differences between description- and experience-based tasks (Haines, Kvam, & Turner, under review)—both of which we use in the current study.

Under the updating scheme described by Equation 9, the different outcomes *x_ijt_* can be thought of as features that are either observed or not after each choice, similar to category learning models (e.g., Turner, 2019). When they are observed (or not observed), their experience weights increase toward 1 (or decrease toward 0) in proportion to the learning rate and the prediction error. Therefore, experience weights converge toward the probability of observing each outcome, but they fluctuate trial-to-trial based on the recent history of outcomes. Note that although experience weights converge toward outcome probabilities, they are not proper probabilities as they do not sum to 1 (akin to the “decision weights” in prospect theory; Kahneman & Tversky, 1979). For intuition, imagine a participant making a choice between action A (“$3 with Pr(1)”) and action B (“$4 with Pr(.8), 0 otherwise”). If they choose A and are subsequently given feedback that A returned $3 and B returned $4, they increase experience weights corresponding to $3 and $4 and decrease the experience weight of $0. An important implication of this updating process is that a given counterfactual experience (e.g., receiving $3 when the alternative choice results in $4) does not need to be fully experienced in order for it to be expected to occur in the future—instead, the experience weights of individual outcomes determine what possible counterfactual outcomes are expected. In this way, the model can learn to expect regret/disappointment in tasks using partial feedback (i.e. where only the chosen action’s outcome is revealed), which lead to slower changes in behavior relative to full-information feedback tasks (e.g., Rakow et al., 2015).

In the context of the task we used, feedback is not given for the first 5 trials, so there is no updating and all experience weights therefore remain at 0. Starting from the 6^th^ trial, the outcomes are shown for both chosen and unchosen options after each choice. Therefore, using Equation 9, we assume that changes in choice behavior starting from the 6^th^ trial result from the effects of participants learning to represent the probability of counterfactual outcomes over time as they experience each outcome. This counterfactual learning process is similar in kind to reversal/fictive learning models (e.g., Gläscher, Hampton, & O’Doherty, 2009; Lohrenz, McCabe, Camerer, & Montague, 2007), although our proposed mechanism is more general due to its explicit representation of outcome probabilities which allows for a much closer correspondence to classic regret/disappointment theory.

#### Disappointment Minimization Learning Model

The Disappointment Minimization Learning model assumes that subjective expected pleasure (*SEP*) for each choice option *i* on trial *t* is given by:

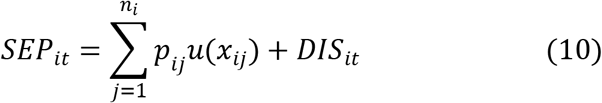

where the terms in the summation are the same as described in the previous model (see *p_ij_* and *u*(*x*) in Equations 4-5), and *DIS_it_* is a trial-dependent disappointment expectation that is determined according to the following delta updating rule:

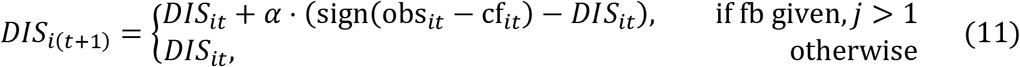

Here, we formalize disappointment as the sign of the difference between the observed outcome for each option (obs_*it*_) and the alternative, counterfactual outcomes (cf_*it*_) that could have occurred if the choice turned out differently. For example, if option A can return either −$4 or −$12, and −$4 is observed on the current trial, disappointment for option A is equal to sign(obs_*it*_ − cf_*it*_) = sign([−4] − [−12]) = 1 and *DIS_it_* then increases toward 1 in proportion to the learning rate *α* (indicating greater expectation for *elation*) and prediction error. Similar to the experience weights in the Counterfactual Representation Learning model, we initialize all the expectations to 0 at the start of each game (*DIS*_*i*1_ = 0). Further, *SEP* values are entered into the same softmax choice rule described by Equation 8, along with the persons-specific choice sensitivity parameter (*θ*).

Equation 11 produces an increasing preference for options that tend to return their highest outcome (i.e. the best outcome within an option), thus minimizing the probability of experiencing disappointment. Note that updating only occurs for a given action *i* if it can return more than 1 potential outcome (i.e. *j* > 1) and if feedback (fb) is presented on the current trial, indicating that disappointment cannot occur when an outcome is guaranteed. This behavior is similar to other reinforcement learning models that track the expected win frequency of each action, which leads to an increased preference for actions that win most frequently (see Equations 7-9 from Haines, Vassileva, & Ahn, 2018). It also produces behavior consistent with win-stay-lose-shift heuristics (e.g., Worthy, Hawthorne, & Otto, 2013).

#### Regret Minimization Learning Model

The Regret Minimization Learning model assumes that *SEP* is given by:

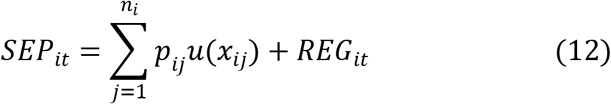

where *u*(*x*) is the same utility function as in previous models, and *REG_it_* is a trial-dependent regret expectation that is similar *DIS_it_*, but is computed *across* options as opposed to *within* options:

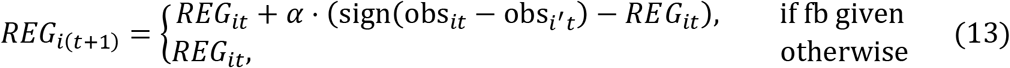

Here, we formalize regret as the sign of the difference between the observed outcomes across options. Like the models described above, we initialize all regret expectations to 0 at the start of each game (*REG*_*i*1_ = 0), and we use the softmax choice rule with choice sensitivity parameter (Equation 8) to transform *SEP* into choice probabilities. Therefore, Equation 13 produces learning dynamics that lead to an increasing preference for the action that most frequently returns the best outcome across actions on a trial-to-trial basis, thus minimizing the probability of experiencing regret. Such behavior is similar to models including the Best Estimate and Sampling Tools model, which minimizes immediate regret through a sampling mechanism that recalls the outcomes of each option on a past trial and prefers the one with the best outcome (see Equation 1 from Erev et al., 2017). Equation 13 differs in that it learns a regret expectation (i.e. it integrates over the recent history of trials) as opposed to relying on a single previously experienced trial.

#### Disappointment/Regret Minimization Learning Model

The Disappointment/Regret Minimization Learning model combines the Disappointment and Regret Minimization models described above, such that *SEP* is given by the subjective utility of each option plus the trial-dependent *DIS* and *REG* expectations. Therefore, it is most similar to the Counterfactual Representation Learning model, in that both disappointment and regret expectations are assumed to develop with experience as outcomes are observed. However, it is different from the Counterfactual Representation Learning model in that it does not explicitly represent each outcome probability to compute counterfactual expectations—instead, expectations are the recency-weighted average of past realized counterfactual comparisons (see Equations 11 and 13).

#### Expectation Maximization Learning Model

The Expectation Maximization Learning model functioned as a baseline model for the current study. We included it to determine if the counterfactual disappointment and regret learning models offered a better account of changes in participants’ preferences with experience relative to simple recency effects where each option and outcome is valued independently. Specifically, *SEP* is given by:

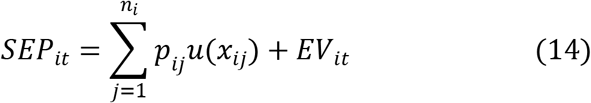

where *EV_it_* is a trial-dependent expected value term that is updated using the following delta rule:

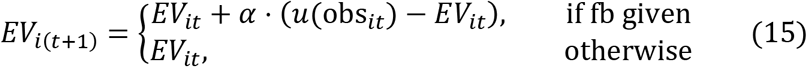

Here, *u*(*x*) is the same utility function as described in previous models, and *EV_it_* is updated toward the utility of the observed outcome for each option *i*, which produces increases in preference for the option that shows a higher average subjective value across recent trials. We use the same softmax choice rule described by Equation 8 to transform *SEP* into choice probabilities. Therefore, the Expectation Maximization Learning model is similar to models such as the naïve sampler model and other basic reinforcement learning models (see Erev & Roth, 2014), which tend to prefer options that return the best average payoff within a small sample of recently experienced outcomes.

### Model fitting procedure

We fit all models using hierarchical Bayesian analysis (HBA), with separate hierarchical models fit to each of the 7 sets of games. We chose to fit separate hierarchies to each set for mostly computational purposes^1^, but also because each set consisted of different participants and games. HBA allows for individual-level (i.e. participant-level) parameter estimation while simultaneously pooling information across participants to increase certainty in individual-level estimates. Further, HBA has previously been shown to provide better parameter recovery than traditional methods such as individual-level maximum likelihood estimation (MLE) (e.g., Ahn, Krawitz, Kim, Busemeyer, & Brown, 2011), suggesting that individual-level HBA estimates can be interpreted with more confidence compared to traditional MLE estimates. We used Stan (version 2.15.1) to implement HBA. Stan is a probabilistic programming language that employs the No-U-Turn Hamiltonian Monte Carlo (HMC) sampler, which is a variant of Markov Chain Monte Carlo (MCMC), to efficiently sample from the joint posterior distribution across all specified model parameters (Carpenter et al., 2017).

We used a standard convention to parameterize the prior distributions for all model parameters (Ahn, Haines, & Zhang, 2017). Specifically, we assumed that each set of individual-level parameters was drawn from a group-level distribution. We assumed normal group-level distributions, where prior means and standard deviations were set to normal distributions. We used non-centered parameterizations to decrease the dependence between group-level mean and standard deviation parameters (Betancourt & Girolami, 2013). Bounded parameters (e.g., *α* ∈ (0, 1)) were estimated in an unconstrained space and then inverse Probit-transformed (i.e. the cumulative distribution function of the standard normal) to the constrained space to maximize MCMC sampler efficiency (Ahn et al., 2014; 2017; Wetzels, Vandekerckhove, Tuerlinckx, & Wagenmakers, 2010). Once transformed to the constrained space, parameters with upper bounds greater than 1 were scaled accordingly. For example, the learning rate parameter for each of the models was parameterized as:

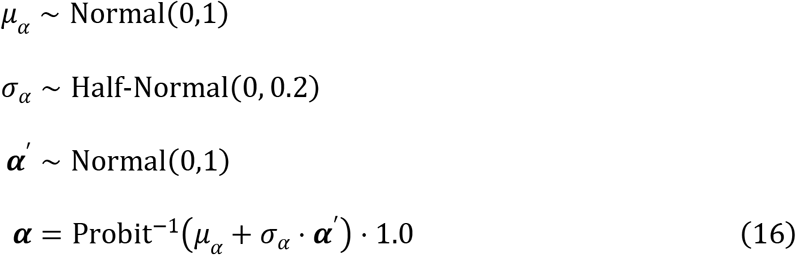

Here, *μ_α_* (−∞ < *μ_α_* < +∞) and *σ_α_* (0 < *σ_α_* < +∞) are the group-level mean and standard deviation, respectively, and bold terms indicate vectors of individual-level parameters. This parameterization assumes that the prior distribution over each individual-level parameter is near-uniform, and that variance is relatively low across participants (which provides regularization of extreme individual-level deviations toward the group-level mean). Since 0 < *α* < 1, the scaling factor was set to 1.0 in the example above. We used the same non-centered parameterization for *ω* and *c* (including the same prior distributions), except they were scaled by 1.5 and 5.0, respectively.

We ran all models for 1,500 iterations across 4 separate sampling chains, with the first 500 samples as warm-up (analogous to burn-in in Gibbs samplers) for a total of 4,000 posterior samples for each parameter. For all models, we checked convergence to the target joint posterior distribution by visually inspecting trace-plots and ensuring that all Gelman-Rubin (a.k.a. 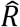) statistics were below 1.1, which suggests that the variance between chains is lower than the variance within chains (Gelman & Rubin, 1992). R and Stan codes for the cognitive models will be made available on the *hBayesDM* package (Ahn et al., 2017) upon publication.

### Model comparison

We used two different methods to compare models, namely: (1) penalized model fit, which is a statistical measure of how accurately a model can predict participants’ choices on the next trial given their fitted model parameters, choice history, and a penalty term for model complexity; and posterior predictive simulations, which involve graphically comparing the difference between participants’ entire choice histories and choice histories simulated from their fitted model parameters to determine which model best accounts for changes in behavior across time. We used multiple methods because previous studies consistently show that different model comparison methods can lead to different conclusions, and because fit statistics alone (e.g., mean squared error, AIC, BIC, etc.) cannot ascertain which model can best “explain” observed data (Ahn et al., 2008; 2014; Haines, Vassileva, & Ahn, 2018; Navarro, 2019; Steingroever et al., 2014; Yechiam & Ert, 2007). Therefore, in judging model performance, we use statistical fit and qualitative visual checks to guide our assessment, but also interpret the model holistically in terms of how plausible the implied cognitive mechanisms are.

#### Penalized model fit

We assessed penalized model fit using the leave-one-out information criterion (LOOIC; Vehtari, Gelman, & Gabry, 2016), which is a fully-Bayesian analogue of traditional, commonly used information criteria (e.g., AIC and BIC). LOOIC approximates true leave-one-out prediction accuracy, and it can be computed using the log pointwise posterior predictive density (LPPD) of a fitted model. To compute the LPPD of each competing model, we calculated the log likelihood of each participants’ true choice on trial *t* + 1 conditional on their parameter estimates and choice history (i.e. trials ∈ {1,2,…, *t*}). Therefore, LOOIC compliments the posterior predictive simulations described above, as it provides a quantitative measure of how well each model can capture participants’ trial-to-trial choices while accounting for parameter uncertainty. The log likelihood was calculated for each posterior sample and summed within each participant across games (preserving all posterior samples), resulting in an *S* × *N* LPPD matrix where *S* and *N* are the numbers of posterior samples and participants, respectively. We used the *loo* R package, which is developed by the Stan team (Vehtari et al., 2016), to compute LOOIC values from the LPPD matrix. Note that LOOIC is on the deviance scale where lower values indicate better model fit (including complexity penalization).

#### Posterior predictive simulations

Posterior predictive simulations are similar to the post-hoc absolute fit measure used in previous studies (e.g., Steingroever et al., 2014; Steingroever, Wetzels, & Wagenmakers, 2013), but they differ in that they convey uncertainty in simulated choices that is attributable to the underlying parameter estimates. Further, they differ from methods such as the absolute simulation method, which uses estimated parameters to simulate behavior on the task without conditioning on participants’ actual trial-level choices (e.g., Haines, Vassileva, & Ahn, 2018; Steingroever et al., 2014; Steingroever, Wetzels, & Wagenmakers, 2013). We generated posterior predictions by first fitting each model to participants’ choice data, followed by simulating participants’ trial-to-trial choices using the full joint posterior distribution of their fitted model parameters and by conditioning on their actual trial-level choices. Note that we fit a different hierarchical model to each set of games, but parameters were fixed within-participants across games within each set. We then averaged across participants and compared the posterior predictive simulations and the true across-participant average choice proportions for each of the 105 games using graphical measures. Our use of posterior predictive simulations allows for us to determine if each model can capture changes in participants’ observed behavior in response to the feedback that they start to experience on the 6^th^ trial of each game, which could be obscured if we used fit statistics alone to conduct model comparison. For example, a model could exhibit relatively good fit statistics but fail to capture learning effects observed in choice rates across trials. Posterior predictive simulations therefore allowed us to identify specific areas of model misfit and determine whether models captured the effects of feedback on changes in behavior across trials.

## Study 1 Results

### Model comparison: Penalized model fit

The Counterfactual Representation and Regret Minimization Learning models showed the best penalized model fit statistics across all 7 sets, and the Counterfactual Representation Learning model performed best overall (see Figure 3). These results corroborate our posterior predictive simulations.

**Figure 3.**
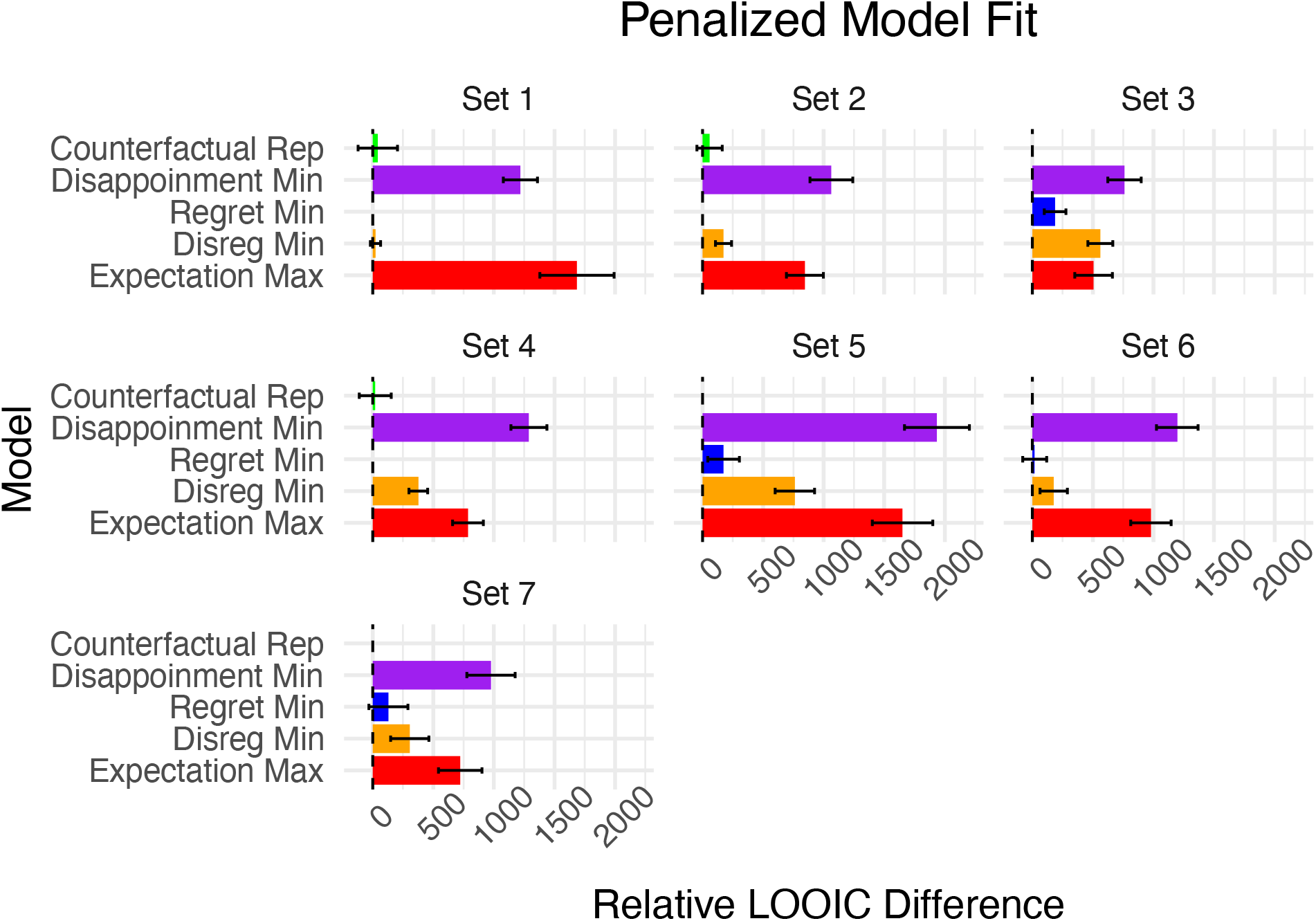
Penalized model fit in the context of description-based games. Leave-one-out information criterion (LOOIC) scores relative to the best fitting model within each set of games. Lower LOOIC values indicate better model fit accounting for model complexity. Error bars represent ± 1 standard error of the difference between the best fitting model in the set and respective competing models.

### Model comparison: Posterior predictive simulations

Figure 4 shows both the true choice rates and posterior predictive simulations averaged across participants for 6 representative games within set 5. We only show these particular games for brevity, but we summarize our observations across all sets and games here (see Figure’s S1-20 in the Supplementary Text for all 105 games). Overall, the Counterfactual Representation Learning, Regret Minimization Learning, and Disappointment/Regret Minimization Learning models best captured changes in participants’ behavior across trials, indicating that regret expectations play a crucial role in facilitating changes in behavior in response to full information feedback. Conversely, the Disappointment Minimization and Expectation Maximization models poorly tracked the observed choice proportions across trials, and in fact often showed little to no changes in predictions in response to feedback across games despite participants showing strong changes in preferences. Of the models containing regret terms, across all games, the Counterfactual Representation Learning model more rapidly converged toward participants’ observed changes in choice behavior, indicating that people may explicitly represent and update counterfactual outcome probabilities (i.e. the counterfactual “experience weight” in Equation 9) as opposed to tracking a running expectation of regret and disappointment. Explicit representation of counterfactual outcomes and probabilities is consistent with traditional models of regret and disappointment (Loomes & Sugden, 1982; 1986), along with empirical evidence that people make counterfactual comparisons between pairs of different possible outcomes (Ordóñez, Connolly, & Coughlan, 2000).

**Figure 4.**
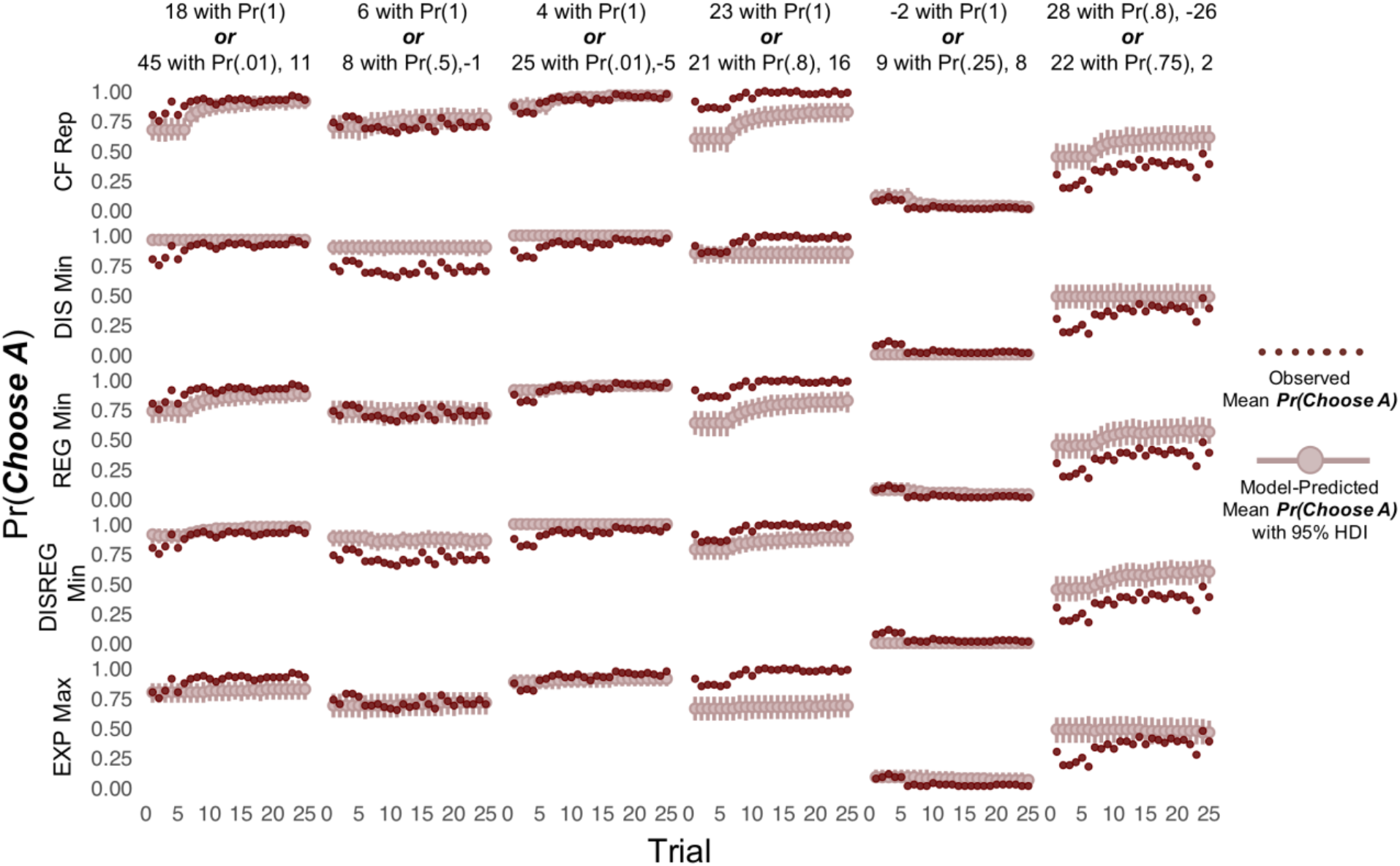
Posterior predictive simulations for a subset of games. Posterior predictive simulations for a representative sample of games from set 5 (see Table S1 for information all sets and games, and Figure’s S1-20 for posterior predictive simulations for all sets and games). Different games are shown across columns (labels describing the game), and the competing models are shown across rows. Here, dark red points indicate the mean choice proportions across participants, and lighter red points with intervals represent the mean of the posterior predictive simulations across participants with 95% highest density intervals (HDI). CF Rep = Counterfactual Representation Learning model; DIS Min = Disappointment Minimization Learning model; REG Min = Regret Minimization Learning model; DISREG Min = Disappointment/Regret Minimization Learning Model; EXP Max = Expectation Maximization Learning model.

Although the models with regret terms performed well for most games, there were some notable exceptions. Specifically, the models tended to underestimate preferences for games wherein one choice option always returned a better outcome than the alternative option. For example, Figure 4 shows a game where participants chose between option A (23 with Pr(1)) or option B (21 with Pr(.8), otherwise 0). Here, option A is always the best choice—it has the highest expected value (albeit by a small amount), and it will never result in either disappointment or regret. As expected, participants almost exclusively choose option A. However, the models all show a much less pronounced preference for option A (although it is still preferred over option B on average). We observed this pattern across many games, all sharing the feature that one option will always return a better outcome than the alternative. These patterns are consistent with the *certainty effect*, wherein certain outcomes (i.e. those with Pr(1)) are given higher “decision weights” relative to uncertain outcomes (e.g., Allais, 1953). Importantly, the certainty effect can be accounted for using probability weighting functions such as those proposed by prospect theory (Kahneman & Tversky, 1979). Therefore, although we did not include a probability weighting function in our models, such an extension could potentially allow the models to better capture certainty effects (e.g., replacing *p_ij_* in Equation 4 with *f*(*p_ij_*), where *f*(*x*) is a non-linear probability weighting function).

## Interim Discussion

Our first study revealed that participants’ changes in behavior in response to full information feedback during repeated description-based decisions are best explained by models assuming that participants make efforts to minimize expected regret. Overall, the best performing model assumed that people learn to represent the probability of counterfactual events (i.e. “experience weights” for disappointment, regret, and their counterparts elation and rejoice) as they gain experience with each possible outcome. This “feature-based” account of counterfactual learning (where different numerical outcomes are conceptualized as different features that are observed or not after a choice) contrasts the competing “running expectation-based” models that we tested, all which assume that counterfactual events are integrated into a single value signal for each choice option that is learned with experience. These results show that pairwise difference formulations of disappointment and regret theory are able to account for changes in behavior over time as a result of experience.

Although findings from our first study offered insight into cognitive mechanisms that may underly counterfactual thinking, the models did not incorporate experienced emotion, which limits our ability to test functional theories of counterfactual thinking as described in the introduction. Further, the task from study 1 gave participants complete information on the probabilities of various outcomes for each choice. Therefore, in study 2 we: (1) determine whether the models tested in study 1 can capture pure experience-based decisions (where probabilities and outcomes are not given and must be learned), and (2) identify how participants’ real-time emotional experiences interact with assumed cognitive mechanisms to facilitate goal-oriented changes in behavior.

## Study 2 Method

### Participants

Our second study included 51 participants’ data collected from two different research sites. Of these participants, 31 had facial expressions recorded, while 19 are from a previous study which did not record facial expressions (Ahn et al., 2012). We aggregated both datasets for the purposes of the current study. All participants gave informed consent prior to participating in the study. Our study protocol was approved by The Ohio State University’s Institutional Review Board (protocol #2016H0108).

### Behavioral task

All participants completed four separate gambling games in randomized order. Participants were told that each game was independent of all other games, and they were given an opportunity for a break between each game. Each game consisted of 90 trials, where participants were asked to choose between two options (see Figure 5). Throughout each game, selecting one of the options won a fixed amount of points (i.e. safe option), whereas the other option had some probability of winning a high or low amount of points (i.e. risky option). Locations of the safe and risky options were randomized across participants but remained fixed within games. The probability of winning a high number of points for the risky option was fixed but unknown, and participants had to learn the probability from experience. After making a choice, point values for both the chosen and unchosen options were revealed (i.e. “full-information feedback”), which allowed participants to make counterfactual comparisons between the choices they made and could have made. Participants were instructed to make choices that maximized their points. Unbeknownst to participants, the expected value for each option was identical (see Table 1 for payoff distributions for each game).

**Figure 5.**
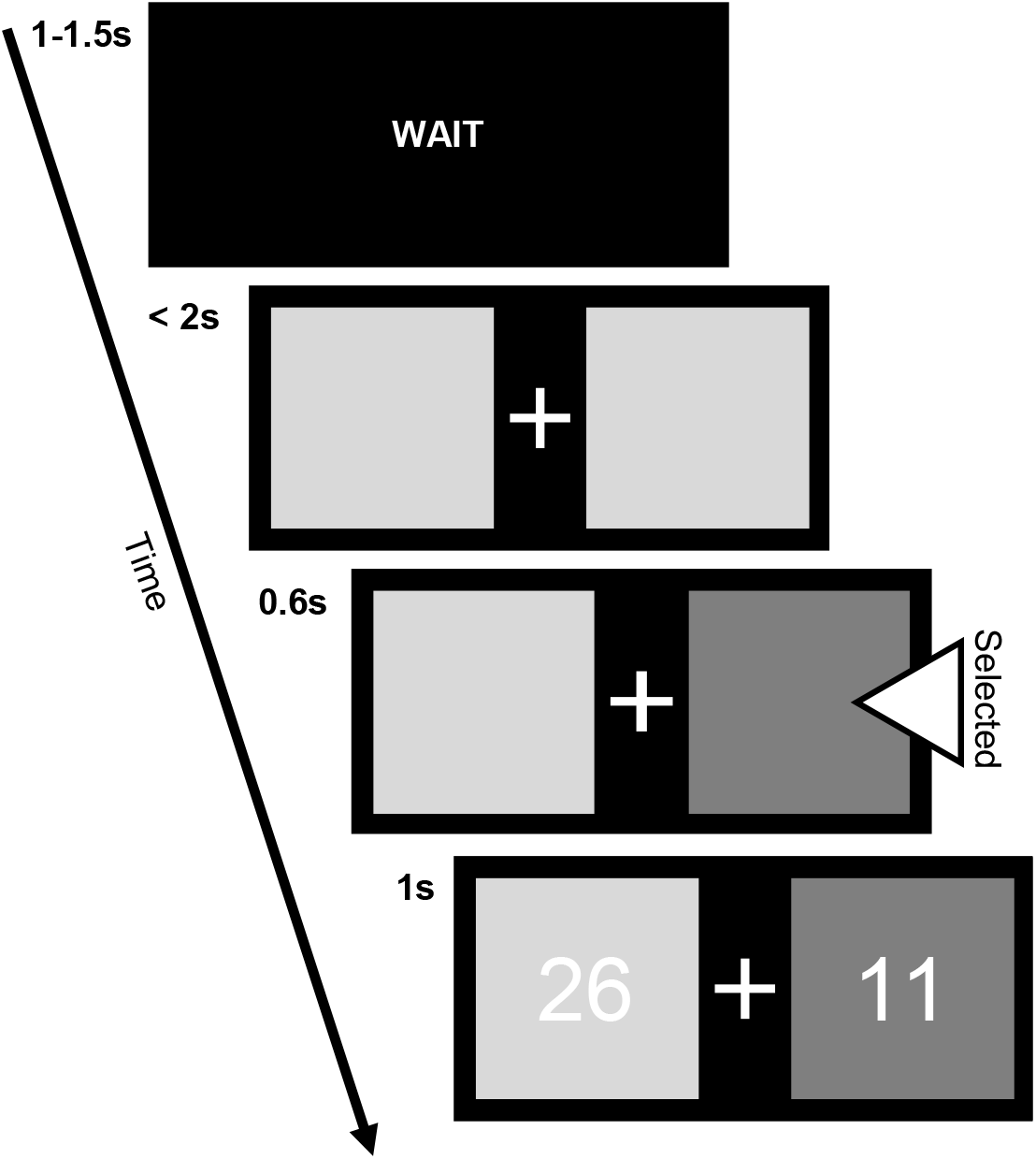
Pure experience-based behavioral task with full-information feedback. Unlike study 1, the task from study 2 did not describe the outcomes and associated probabilities of each choice to participants. However, the tasks are similar in that they both offer full-information feedback after a choice is selected.

**Table 1.** Payoff structure for each game in Study 2

| Game | Option A (safe) |  |  |  | Option B (risky) |  |  |  |
| --- | --- | --- | --- | --- | --- | --- | --- | --- |
| | $A_H$ | (Pr<sub>H</sub>) | $A_L$ | (Pr<sub>L</sub>) | $B_H$ | (Pr<sub>H</sub>) | $B_L$ | (Pr<sub>L</sub>) |
| 1 | 12 | (1) | – | – | 56 | (.2) | 1 | (.8) |
| 2 | 11 | (1) | – | – | 26 | (.4) | 1 | (.6) |
| 3 | 10 | (1) | – | – | 16 | (.6) | 1 | (.4) |
| 4 | 9 | (1) | – | – | 11 | (.8) | 1 | (.2) |

*Note.* $A_H$ = high outcome for option A. $A_L$ = low outcome for option A (same notation for option B). Pr<sub>H</sub> = high outcome for given option. Pr<sub>L</sub> = low outcome for given option. Note that outcomes and their corresponding probabilities were not described to participants. Instead, participants had to learn from experience-based feedback.

### Experience-based computational models

We applied the same five models described in study 1 to data collected from the pure experience-based paradigm used in study 2 (e.g., Figure 5). However, because the models from Study 1 assume that participants have access to the outcome values and probabilities of each choice option before receiving any feedback (e.g., Figure 2), we made a simple modification to each models to reflect the absence of described outcome information in the experience-based paradigm. We describe these modifications below.

For the Counterfactual Representation Learning model, the described probabilities only appear in Equation 4, wherein they function to weight the modified utility term. Importantly, because the model explicitly represents probabilities in the form of “experience weights” (i.e. ***W*** in Equation 9), the described probability from Equation 4 can simply be replaced with the subjective experience weight to extend the model to pure experience-based decision-making:

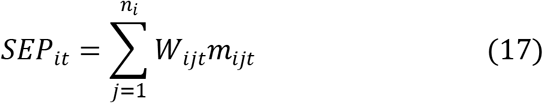

Here, *W_ijt_* is the experience weight for the corresponding outcome *x_ij_*. All other aspects of the model are identical to the description-based version (including the updating rule and initializing all ***W*** = 0). Therefore, whereas the description-based version reduces to a standard prospect theory like model before any feedback, the experience-based version has a uniform preference for all options before feedback because all experience weights start at 0 (and *SEP* is thus 0 for all options).

For the remaining models, since they do not explicitly represent the probability of each outcome occurring, we dropped the terms containing the described probability (*p_ij_*) from each model. Using the Disappointment Minimization Learning model as an example, we dropped the summation term from Equation 10, such that *SEP_it_* = *DIS_it_*. We did the same for all other models. Therefore, each model assumes that *SEP* is given by a recency weighted average over past counterfactual experiences (except for the Expectation Maximization Learning model, which assumes that the *SEP* of each option is given by a recency weighted average of observed outcomes for that option).

### Model-based facial expression analysis

#### Automated facial expression coding

To measure the valence of participants’ facial expressions during feedback, we used an automated facial expression coding (AFEC) model that we developed in a previous study (Haines et al., 2019). The AFEC model was trained to code for positive and negative affect intensity on a scale from 1 (*no affect*) to 7 (*extreme affect*), where positive and negative affect are coded separately rather than on a polarized positive–negative valence continuum. The AFEC model first uses FACET—a computer vision software (iMotions, 2018)—to detect the presence of 20 different facial action units (Ekman, Friesen, & Hager, 2002), which are then translated to affect intensity ratings using a machine learning model that we previously validated. In our validation study, the model showed correlations with human observer ratings of .89 and .76 for positive and negative affect intensity, respectively (for more details, see Haines et al., 2019).

To preprocess and apply the AFEC model to our participants’ facial expressions in response to outcome feedback, we followed four steps. First, we used FACET to detect the presence of 20 different facial action units (AUs) during the feedback phase of the task. FACET-detected AUs are derived from the anatomically-based Facial Action Coding System (FACS; Ekman et al., 2002), which is a widely-used facial coding systems. FACET outputs a time-series of values for each AU at a rate of 30 Hz, where values represent the probability (i.e. “evidence”) that a given AU is present in each sample. Second, we computed the area-under-the-curve (AUC) of each AU time-series and divided (a.k.a. normalized) the resulting value by the total length of time that a face was detected throughout the 1 second feedback phase in the task (per trial). Normalization ensures that clips of varying quality (e.g., 70% versus 90% face detection accuracy) do not affect the magnitude of the AUC values, which is important for the machine learning step. We excluded any trials where a participant’s face was detected for less than 10% of the total 30 samples in the given trial (~3% of trials excluded in total). Third, we iterated step 2 for each trial and participant. Fourth, we entered the resulting values as predictors in the AFEC model described above to generate valence intensity ratings for positive and negative affect (Haines et al., 2019). We used the positive and negative affect ratings as input into the computational models as described below.

#### Incorporating facial expressions into computational models

To determine whether emotional facial expressions reflected choice mechanisms of learning (*α*), outcome valuation (*ω*), or exploration/exploitation (*c*), we developed 3 competing models that used the positive and negative affect intensity scores to modulate trial-by-trial model parameters (*α*, *ω*, *c*). To do so, we computed an overall valence score for each trial by taking the difference in positive and negative affect ratings for the given participant and trial. We chose this approach over modeling positive and negative affect as separate dimensions to reduce the number of possible models for model comparison purposes. We then standardized (i.e. z-scored) the valence ratings across participants, games, and trials before using them as input to the model. We parameterized each model such that the respective model parameter for trial *t* was a linear combination of a baseline parameter and a parameter determining the effect of emotion valence intensity. For example, the learning rate parameter was determined by:

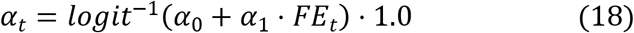

Here, *α*_0_ and *α*_1_ indicate the baseline learning rate and effect of facial expression valence intensity on the learning rate for current trial (*α_t_*), respectively, and *FE_t_* is the standardized facial expression valence rating on trial *t*. Note that the inverse logit function transforms the output so that 0 < *α_t_* < 1, which are the appropriate lower and upper bounds, so we scaled the output by 1.0 (i.e. no scaling). We used the same parameterization for *ω* and *c*, except the scaling factors were 1.5 and 5.0, respectively. On trials where participants’ faces were detected for less than 10% of the feedback stage, we assumed that parameters were not affected by facial expressions. Using the learning rate model as an example, if facial expression data on trial *t* was discarded, then *α_t_* = *logit*^−1^(*α*_0_) ⋅ 1.0. In summary, each of the models makes an explicit assumption about which of the three cognitive mechanisms (i.e. learning, outcome valuation, or exploration/exploitation) is related to moment-to-moment emotional valence intensity, which allowed us to take a model-based approach to explore competing models of behavioral change.

### Model fitting procedure

For the models incorporating facial expressions, we used an alternative prior parameterization for the parameter of interest to reflect the changes described in Equation 18. Using the model assuming that facial expression intensity reflected changes in the learning rate as an example, priors on the modified learning rates were set as follows:

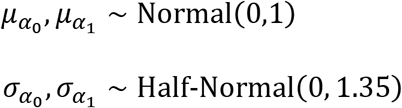

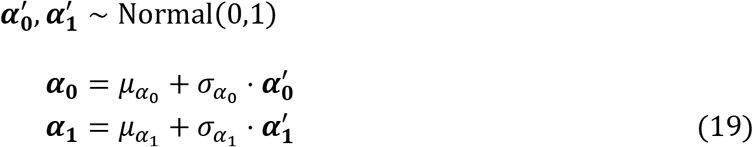

We used these priors because they led to near-uniform priors over the individual-level trial-by-trial parameters (i.e. *α_t_*) after being determined by Equation 18. The same parameterization was used for *ω* and *c* in the models assuming relations between facial expression intensity and reward and choice sensitivity, respectively.

We ran all models for 2,500 iterations across 4 separate sampling chains, with the first 500 samples as warm-up for a total of 8,000 posterior samples for each parameter. We checked convergence to the target joint posterior distributions by visually inspecting trace-plots and ensuring that all Gelman-Rubin (a.k.a. 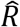) statistics were below 1.1, which suggests that the variance between chains is lower than the variance within chains (Gelman & Rubin, 1992). R and Stan codes for the computational models will be made available on the *hBayesDM* package (Ahn et al., 2017) upon publication.

### Model comparison

We used the same model comparison procedures as described in Study 1 to determine which model performed best in Study 2. However, because we only had facial expression recordings for 31 of the 51 participants, model comparison proceeded in two stages. First, we fit each model to all 51 participants and performed both LOOIC model comparison and posterior predictive simulations to determine whether the findings from Study 1 generalized to Study 2 wherein we used a pure experience-based task. Next, we parameterized the best performing model (the Counterfactual Representation Learning model) using the scheme described in Study 1, and we then fit each model to the subset of 31 participants to identify relations between their facial expression intensity in response to feedback and cognitive mechanisms that could influence behavior. We used LOOIC to identify the best performing facial expression model, which we then used to infer how experienced emotion affected subsequent choice behavior.

## Study 2 Results

### Model comparison: Penalized model fit

Corroborating our findings from Study 1, the Counterfactual Representation Learning model outperformed all other models when assessed using LOOIC (see Figure 6), indicating that it provided the best fit to participants’ observed choice data accounting for model complexity.

**Figure 6.**
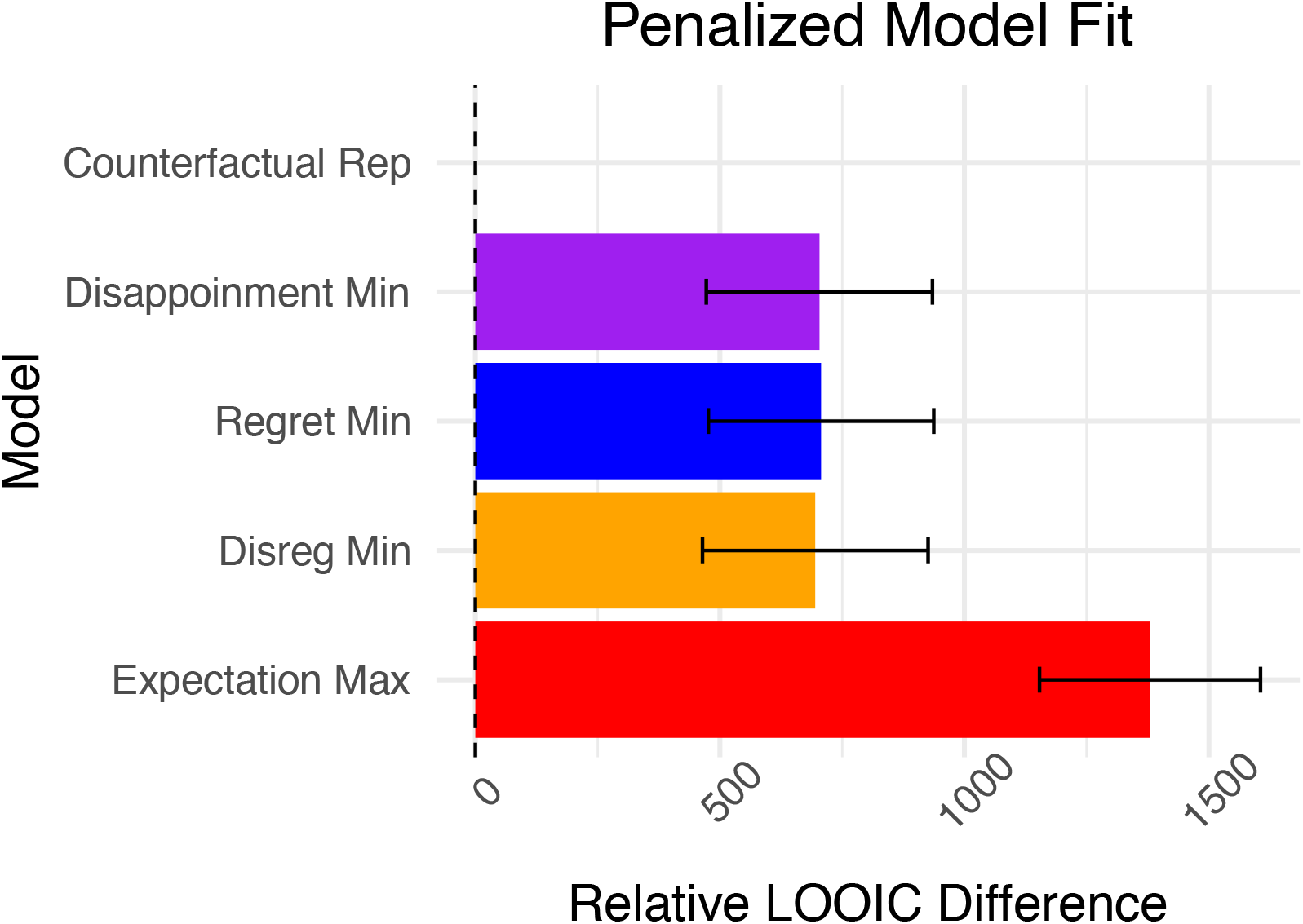
Penalized model fit in the context of experience-based games. Leave-one-out information criterion (LOOIC) scores relative to the best fitting model within across the 51 participants and 4 games from Study 2. Lower LOOIC values indicate better model fit accounting for model complexity. Error bars represent ± 1 standard error of the difference between the best fitting model and respective competing models.

### Model comparison: Posterior predictive simulations

Figure 7 shows both the true and posterior predictive simulations across-participant (*N*=51) choice proportions for the safe versus risky options. Note that we fit each model to all 4 games simultaneously. Despite the safe and risky options having the same expected value within each game, participants showed a clear preference for the risky option in games where the high payoff/extreme outcome was more likely to occur—this pattern of behavior is consistent with previous studies showing that people tend to underweight rare events and/or over-value extreme outcomes when making decisions from experience (e.g., Barron & Erev, 2003; Hertwig et al., 2004; Ludvig & Spetch, 2011; Ludvig, Madan, & Spetch, 2013). Similar to our results from Study 1, the Counterfactual Representation and Regret Minimization Learning models best captured changes in participants’ behavior across trials, suggesting that regret expectations continue to play a crucial role in changes in preference in response to feedback even when outcome distributions are unknown. Further, the Counterfactual Representation Learning model performed slightly better than the Regret Minimization model for game 3, as the Regret Minimization model tended to increasingly prefer option B across trials despite participants not showing such changes (see Figure 7). Overall, our posterior predictions for the pure experience-based task used in Study 2 corroborate our findings from Study 1, suggesting that the explicit representation of counterfactual values and probabilities is general across paradigms.

**Figure 7.**
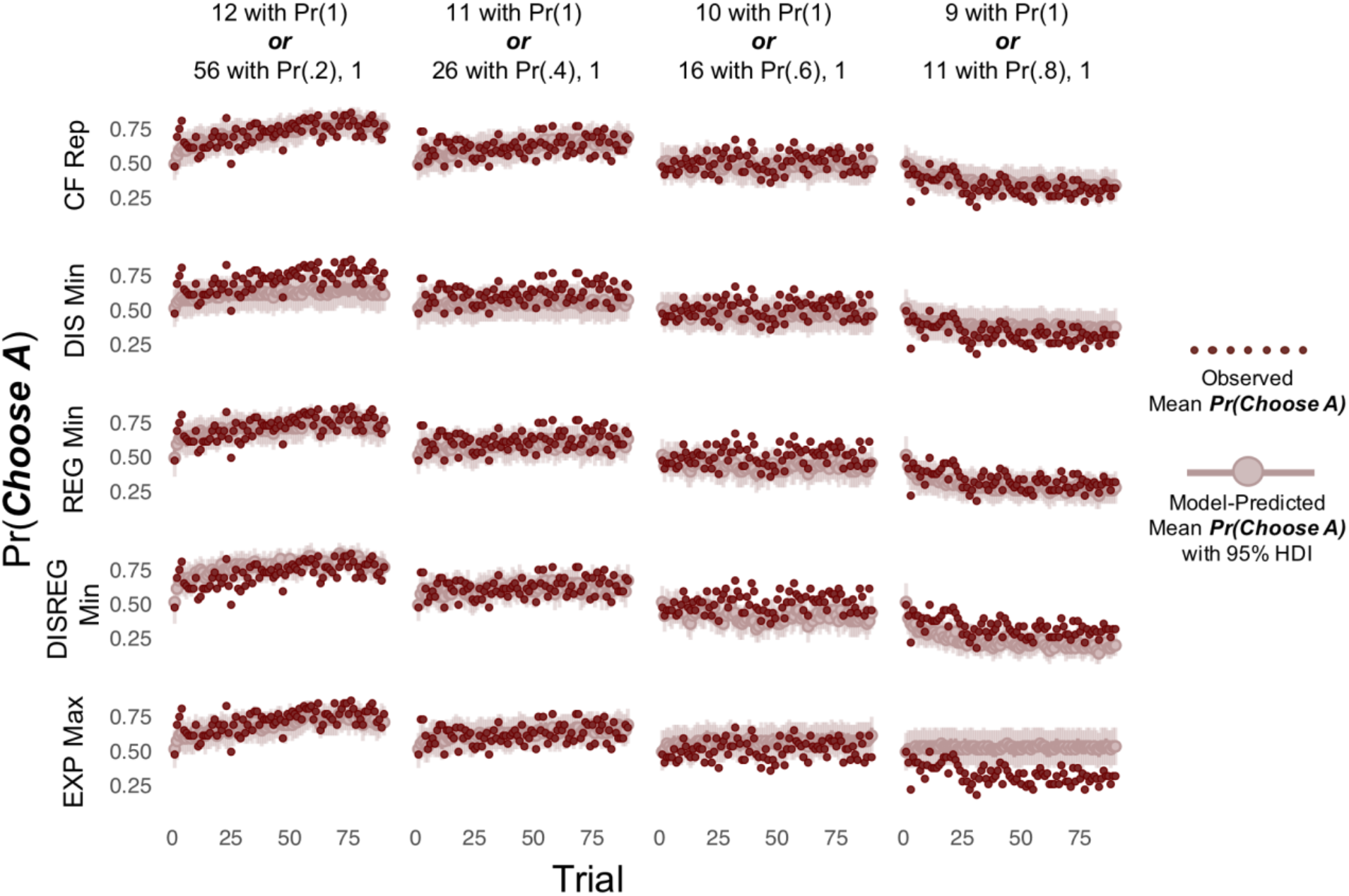
Posterior predictive simulations for all four experience-based games. Posterior predictive simulations for each model across the 4 pure experience-based games in Study 2. Different games are shown across columns (labels describing the game), and the competing models are shown across rows. Here, dark red points indicate the mean choice proportions across participants (*N*=51), and lighter red points with intervals represent the mean of the posterior predictive simulations across participants with 95% highest density intervals (HDI). CF Rep = Counterfactual Representation Learning model; DIS Min = Disappointment Minimization Learning model; REG Min = Regret Minimization Learning model; DISREG Min = Disappointment/Regret Minimization Learning Model; EXP Max = Expectation Maximization Learning model.

### Model-based facial expression analysis

Although our main focus was on overall changes in facial expression valence intensity and not on specific changes in facial expressions, we show a visual depiction of the changes in each of the 20 facial action units (panel A) and overall valence intensity (panel B) in response to different types of feedback in Figure 8 for completeness. In general, the most reliable changes were in facial expressions near the eyes—action units 1, 2, 9, and 43 all exhibited increases in evidence ratings (i.e. an increase in probability of being present) in response to feedback, whereas action units 5 and 7 tended to show decreases. For all these examples, changes were greatest when participants experienced both disappointment and regret on the current trial (e.g., when they chose the risky option and received the low outcome). Activation changed most strongly for AU43, which indicates that participants tended to close their eyes during feedback. This effect is consistent with studies using eye tracking, which show that people often avoid looking at both received and forgone outcomes (Ashby & Rakow, 2016).

**Figure 8.**
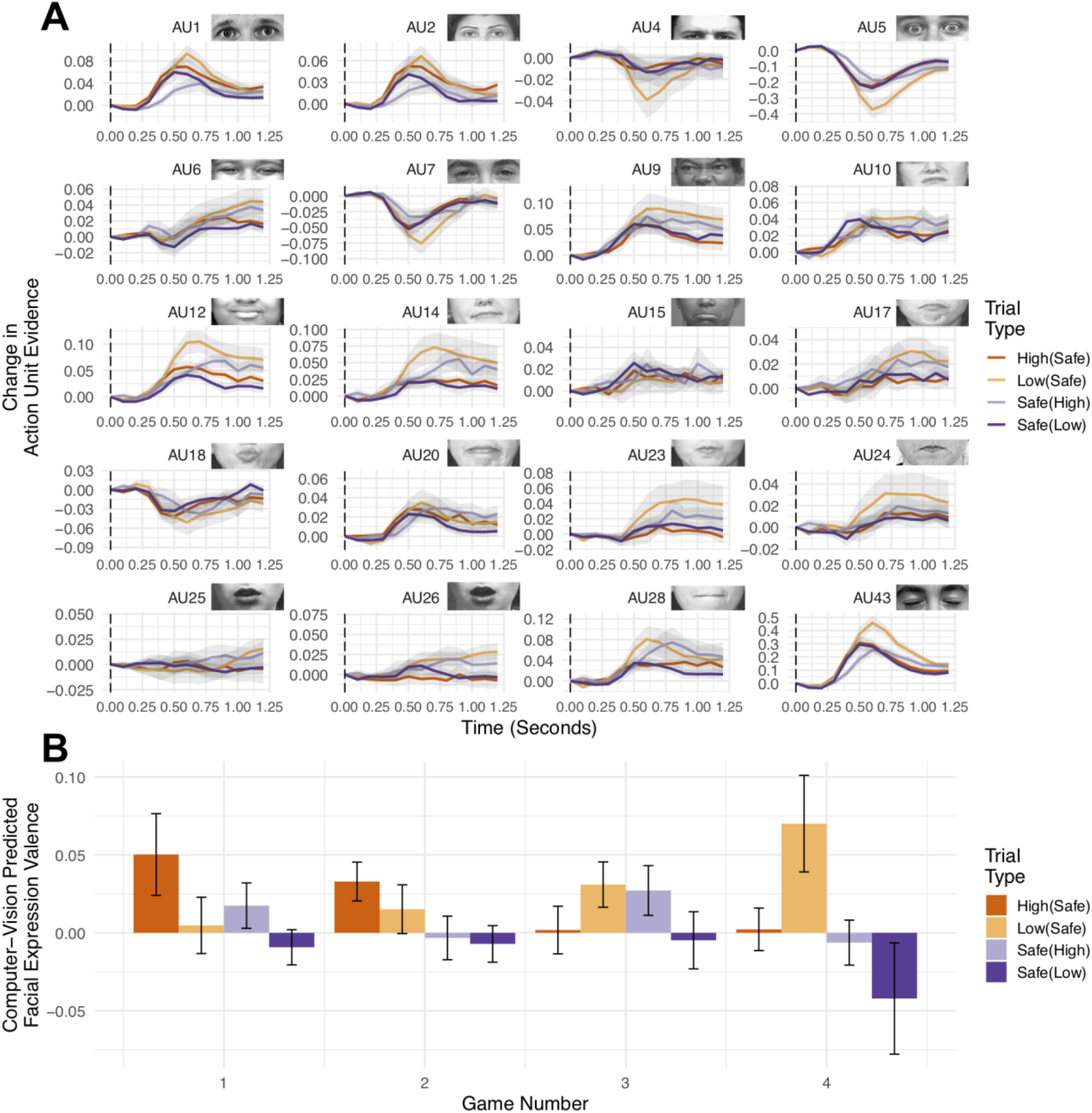
Changes in facial expressions in response to feedback. (A) Changes in computer-vision predicted action unit evidence starting from the moment that feedback is received, separated across the different types of counterfactual outcomes. High(Safe) = chose risky option and received high outcome, foregone safe outcome; Low(Safe) = chose risky option and received low outcome, foregone safe outcome; etc. Change in evidence was computed by first centering the first observation for each trial of the feedback period at 0, and then averaging across trials and participants. Shading indicates ± 1 standard error of the mean across participants. (B) The action unit evidence scores were entered into a secondary model to compute an overall valence score for each trial, and here we show the average changes in overall valence from the choice selection phase (see Figure 4) to the outcome presentation phase across games and different types of outcomes. The uncertainty intervals indicate ± 1 standard error of the mean across participants. Note that these descriptives are only rough proxies for the relationships we aimed to test between facial expression valence and computational model parameters, and we include them here for completeness.

Participants tended to have positive changes in valence in response to receiving the rare outcome upon choosing the risky action across games (i.e. the high outcome in games 1-2 and the low outcome in games 3-4; see Table 1), whereas the most consistent negative change was in game 4 when participants received the safe outcome and the rare, low outcome was foregone. These differences in descriptives across games are somewhat counterintuitive, but they are broadly consistent with regret theory, which predicts that experienced affect is proportional to the expectedness of the outcome. Because expectedness is a function of experience in our task, these descriptives are only rough proxies for the relationships we aimed to test between facial expression valence and computational model parameters. Our next analysis addresses this point, after which we interpret Figure 8B through the lens of the cognitive models.

Because the Counterfactual Representation Learning model showed better overall performance relative to competing models across a variety of games and task paradigms (e.g., description versus pure experience), we used it to further determine how facial expression valence intensity—our primary focus—influenced choice behavior through cognitive mechanisms of learning, reward sensitivity/valuation, or choice sensitivity (exploration/exploitation). Figure 9 shows that the Counterfactual Representation Learning model with a trial-by-trial valence effect on the learning rate parameter, as opposed to outcome valuation or choice sensitivity parameters, provided the best fit, suggesting that the facial expressions that people make in response to feedback relate to how quickly they update their expectations.

**Figure 9.**
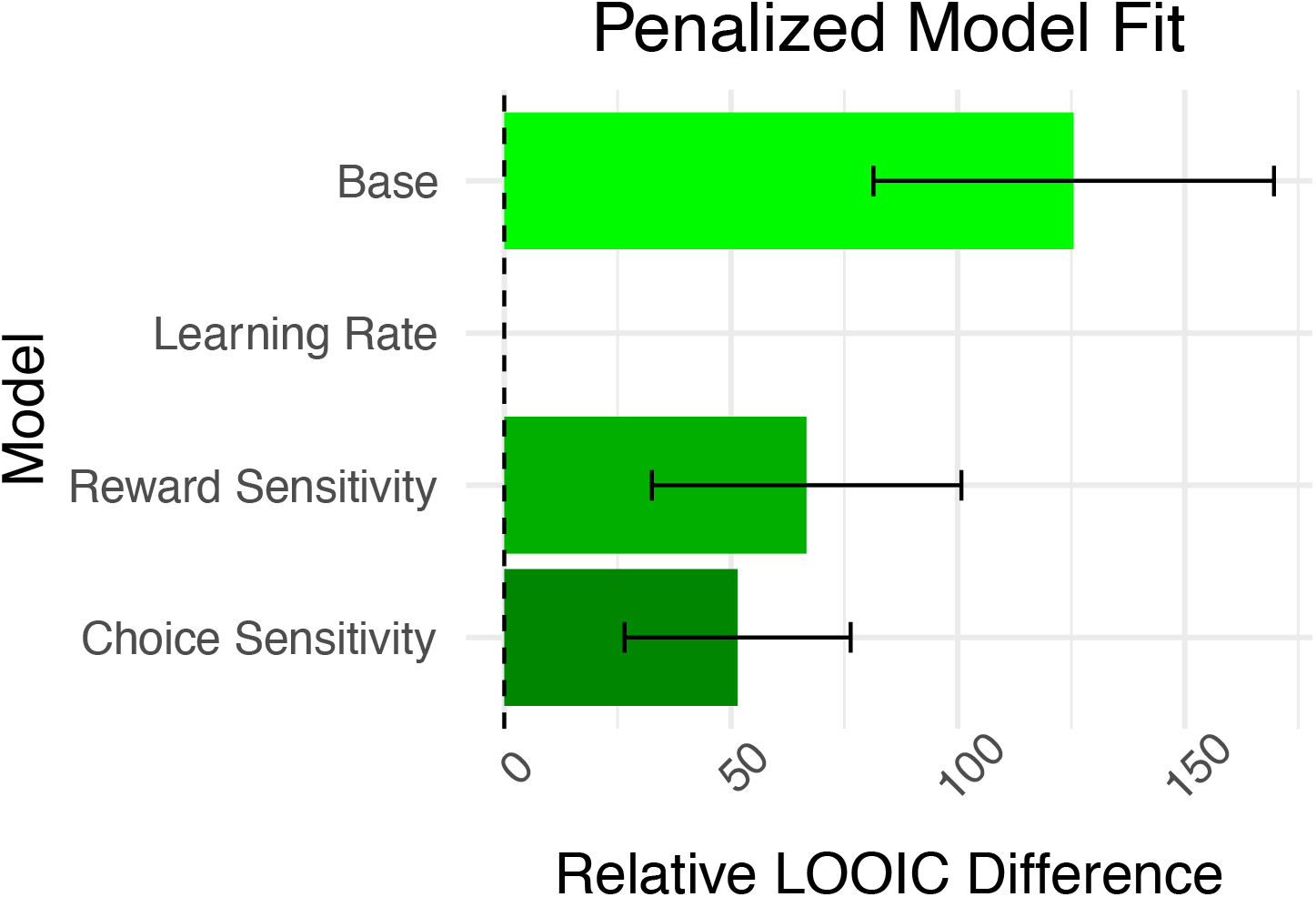
Penalized model fit for models facial expression models. Difference in LOOIC values between models including facial expressions to generate trial-by-trial model parameters. The Base model represents performance of the Counterfactual Representation Learning model for the 31 participants with facial expression recorded, without assuming facial expressions related to model parameters. Learning Rate, Reward Sensitivity, and Choice Sensitivity models assume that trial-by-trial variability in facial expression intensity is associated with changes in *α*, *w*, and *c*, respectively. Error bars represent ± 1 standard error of the difference between the best fitting model and respective competing models.

Figure 10 depicts the effect of facial expression valence intensity on learning. Notably, the group-level posterior distribution of valence on learning 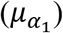 was almost completely negative (95% highest density interval (HDI) = [−1.11, −0.13]), with over 99% of the posterior mass below 0, indicating that participants tend to update their expectations more rapidly as they express more intense negative facial expressions and more slowly as they express more positive facial expressions. This finding is in line with functional theories of counterfactual thinking, which suggest that negative emotions facilitate goal-oriented changes in behavior (Connolly & Zeelenberg, 2002; Zeelenberg & Pieters, 2007). Additionally, it is consistent with previous studies identifying links between surprise and/or prediction error and experienced emotional valence (Mellers, 1997; 1999; Rutledge et al., 2014). Lastly, the relationship between valence intensity and learning rate facilitates interpretation of the descriptive results presented in Figure 8B. When participants made a risky choice and received an unexpected outcome (i.e. the rare outcome), they exhibited positive changes in facial expression valence on average. According to our computational model, these positive (as opposed to negative) changes indicate relatively less expectation updating and subsequently a lower probability if changing choice behavior. This overall pattern is consistent with decision justification theory (e.g., Connolly & Zeelenberg, 2002)—even when participants receive the low outcome in games 3 and 4, the risky choice is “justified” in that the low outcome is rare/unexpected.

**Figure 10.**
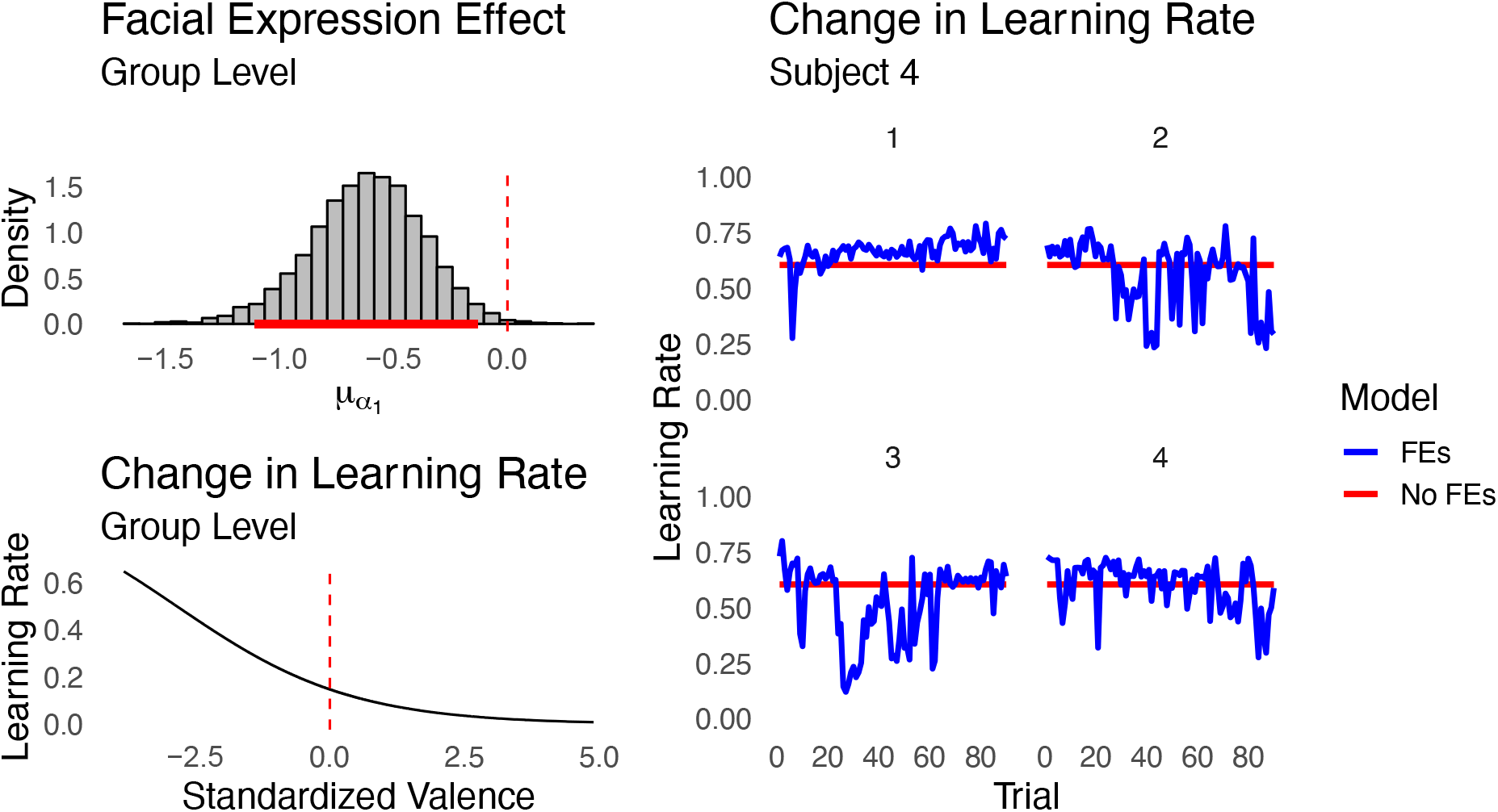
Group- and individual-level effects of emotional valence intensity on learning. Posterior distribution (with 95% highest density interval highlighted in red) over the effect of facial expressions on the learning rate in the Counterfactual Representation Learning model (see Equation 18), a representation of the change in learning rate based on the group-level learning effect 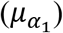 and facial expression valence ratings across participants, and an example from one participant of the learning rate with and without trial-by-trial effects of facial expression valence. Note that as facial expression valence becomes increasingly negative, the learning rate becomes increasingly rapid and vice-versa.

## General Discussion

Our findings are two-fold. First, we found that regret expectations play a key role in changing behavior as people encounter counterfactual information in their decision environment. Moreover, we found that traditional pairwise formulations of regret and disappointment theory can be extended to paradigms wherein people change their behavior with increasing experience. Specifically, the Counterfactual Representational Learning model assumes that people learn to encode their relative “experience weights” with counterfactual outcomes, which correspond to the learned subjective probability of each outcome occurring. Importantly, the explicit representation of outcome probabilities suggested by our model provides insight into functional theories of counterfactual thinking, which predict that people experience more regret when they make poor quality or unjustified decisions (e.g., Inman & Zeelenberg, 2002; Pieters & Zeelenberg, 2005). According to our model, decision quality/justifiability can be thought of in terms of a dynamic experience weight indicating how probable a given regretful or disappointing outcome is in the future. Other studies show that this type of probability estimation can be affected by a number of different processes including recency of experience, similarity to other outcomes, saliency/attention, and differential sensitivity to positive versus negative prediction errors (e.g., Estes, 1976; Haines et al., under review; Lichtenstein, Slovic, Fischhoff, Layman, & Combs, 1978; Zacks & Hasher, 2002). Therefore, to the extent that such processes can be manipulated, we should expect counterfactual expectations to shift over time in predictable ways (e.g., making a regretted outcome more salient should may increase the experience weight for that outcome more so than if it is ignored, etc.). Future studies using the Counterfactual Representation Learning model may benefit from directly manipulating how outcomes are presented or how participants are instructed to evaluate outcomes to determine how to best manipulate counterfactual learning. For example, Sokol-Hessner and colleagues (2009) showed that people are less loss averse when they cognitively reframe losses as “one part of a large portfolio”, rather than “an important decision in isolation”. Such studies may reveal novel insights into decision-making interventions for individuals with psychiatric disorders characterized by emotion dysregulation.

An additional novel prediction made by the Counterfactual Representation Learning model worth exploring in future studies is that counterfactual expectations can influence behavior absent a direct experience with particular joint counterfactual outcomes. For example, if I know that choosing action A can result in either $4 or $0 while action B can result in only $3, Equations 6-7 compute expectations over all possible counterfactual experiences given that I have independently observed each outcome. This account is consistent with traditional models of regret and disappointment (Bell, 1982; 1985; Loomes & Sugden, 1982; 1986), but contrasts contemporary models of experience-based risky decision-making—most which do not assume an explicit representation of values and probabilities and instead compute expectations as averages over past experiences of disappointment and/or regret (e.g., Boorman, Behrens, & Rushworth, 2011; Erev et al., 2014; Hayden, Pearson, & Platt, 2009; Lohrenz, McCabe, Camerer, & Montague, 2007; Yechiam & Rakow, 2012).

Second, our results suggest that people update their counterfactual expectations more rapidly when they experience outcomes while exhibiting intense negative affect (see Figure 10), which may explain why regret and disappointment can lead to negative functional outcomes yet also be looked back upon with appreciation (Kocovski et al., 2005; Lecci et al., 1994; Monroe et al., 2005; Saffrey et al., 2008). For example, negative functional outcomes may be the result of dysfunctional interactions between the cognitive and emotional components of regret that typically facilitate learning and behavioral change, whereas appreciation may result from looking back on regretted decisions that improved later decision-making. Indeed, the orbitofrontal cortex (OFC) and amygdala interact to produce both regret-averse decision-making and extinction learning in healthy adults (Coricelli et al., 2005; Finger, Mitchell, Jones, & Blair, 2008), and altered OFC-amygdala functional/structural connectivity is associated with a number of psychiatric disorders (e.g., Dougherty et al., 2004; Greening & Mitchell, 2015; Hahn et al., 2011; Passamonti et al., 2012). Relatedly, regret appears to have minimal effects on individuals who are highly impulsive and lack trait anxiety (Baskin-Sommers, Stuppy-Sullivan, & Buckholtz, 2016), yet it is overabundant in those with high trait anxiety (e.g, Roese et al., 2009), implicating an important role for regret-driven decision-making in psychological disorders characterized by emotion dysregulation. The rapid updating associated with negative emotion may be useful in fast-changing or uncertain environments, but it could be detrimental when environments are stable (e.g., a consistently high learning rate may lead to an increase in perceived uncertainty despite action-outcome contingencies being consistent). Future studies may use regret-inducing tasks to further explore how the interactions between affective and cognitive components of regret present in individuals with different personality traits and/or psychiatric disorders (see Etkin, Büchel, & Gross, 2015). More broadly, our results are consistent with recent shifts toward conceptualizing emotions as fundamental, adaptive components of human cognition that help us make optimal inferences within constantly changing environments (e.g., Eldar, Rutledge, Dolan, & Niv, 2016).

This is the first study of its kind to include dynamic facial expressions as direct input into a cognitive model, although similar model-based approaches are becoming increasingly common in cognitive neuroscience (Turner, Forstmann, Love, Palmeri, & Van Maanen, 2017). Further, work using automated facial expression coding is gaining traction in social and behavioral sciences due to its efficiency relative to human coders (e.g., Cheong, Brooks, & Chang, 2017; Haines et al., 2019). Future work would benefit from combining automated facial expression coding with behavioral paradigms that collect self-reports of emotion (e.g., Rutledge et al., 2014), which would both allow for more strenuous validity tests of automated measures and create opportunities for exploring the relationships between unobservable and observable emotional states. The advantage of using facial expressions, as opposed to other measurement modalities such as EEG or psychophysiological measures (e.g., heart rate variability, skin conductance, facial electromyography, etc.), is that visual facial features consistently provide the single most predictive measure of reported emotional valence intensity, whereas other modalities are often better suited for arousal (e.g., Chao, Tao, Yang, Li, & Wen, 2015; Kanluan, Grimm, & Kroschel, 2008). We believe that, in the same way that model-based eye-tracking and neuroimaging methods allow for researchers to test the limits of cognitive or computational models with higher fidelity (Turner et al., 2017), facial expression analysis offers another means to testing models that have theoretical implications for emotion, social communication, and related phenomena. Our study offers a proof of concept for the utility of model-based facial expression analysis, and we aim to further validate and extend the general framework in future work.

Although our approach is limited in that we only tested hypotheses regarding valence intensity, future studies may incorporate multiple measurement modalities to better capture different dimensions of emotion. In fact, physiological measures such as skin conductance response, eye-tracking, EEG, and fMRI have previously been used to inform cognitive models (e.g., Cavanagh, Eisenberg, Guitart-Masip, Huys, & Frank, 2013; Frank et al., 2015; Krajbich, Armel, & Rangel, 2010; Jian Li et al., 2011). Moreover, work on joint modeling suggests that model parameters can be estimated more precisely as more measurement modalities are included (Turner, Rodriguez, Norcia, McClure, & Steyvers, 2016). Given the limitations of traditional summary measures of behavioral performance for making inference on individual differences (e.g., Haines et al., 2020; Hedge, Powell, & Sumner, 2017), joint modeling of multiple data modalities may be a fruitful way forward in developing and testing increasingly complex cognitive models. For example, future studies may use the joint modeling approach to explore the social dynamics of decision-making, wherein regret expectations and facial expressions—among other response modalities—play a crucial role in negotiation and trust (Larrick & Boles, 1995; Martinez & Zeelenberg, 2014; Reed, DeScioli, & Pinker, 2014; Reed, Zeglen, & Schmidt, 2012).

Finally, it is worth noting potential extensions to the Counterfactual Representation Learning model. As currently implemented, the probability of each possible outcome (i.e. feature) is learned independently. Although this mechanism works rather well when there are a small number of possible outcomes, it could quickly become inefficient when outcomes are continuous (e.g., if outcomes for each option were drawn from a normal distribution). To remedy this potential shortcoming, the learning mechanism could be modified to include a similarity or attention mechanism, wherein the probabilities of all possible outcomes are upweighted in proportion to their proximity to the observed outcome (e.g., see Turner, 2019). Such learning mechanisms may allow for the model to better generalize to more complex decision contexts than we explored, including multi-stage, multi-attribute, and multi-alternative contexts. Further, although we simplified the utility functions to focus more specifically on learning, it would be worth exploring how differences in utility functions described by traditional models of regret and disappointment (i.e. Equations 1-3) can account for different patterns of decision-making. It is likely that the inclusion of these different utility functions could improve the model’s performance across choice sets.

## Conclusion

Functional theories of counterfactual thinking suggest that the anticipation of positively- or negatively-valenced emotions such as regret/rejoice and disappointment/elation functions to modify behavior in a goal-directed way. Computational models based on principles of regret theory, disappointment theory, and reinforcement learning offer formal mechanisms to explain how people learn to anticipate such cognitive-affective states. Model-based facial expression analysis then provides a principled way of linking the cognitive mechanisms to observed, momentary measures of affect intensity.

## Acknowledgements

We would like to thank both Jerome R. Busemeyer and Brian F. O’Donnell for their insightful comments and suggestions on previous versions of this manuscript. The research was supported by the Basic Science Research Program through the National Research Foundation (NRF) of Korea funded by the Ministry of Science, ICT, & Future Planning (NRF-2018R1C1B3007313 and NRF-2018R1A4A1025891), the Institute for Information & Communications Technology Planning & Evaluation (IITP) grant funded by the Korea government (MSIT) (No. 2019-0-01367, BabyMind), and the Creative-Pioneering Researchers Program through Seoul National University to W.-Y.A.

## Footnotes

1 A single model fit across all 686 participants would require more computer RAM that we have available for making posterior predictions (e.g., participants × games × trials × MCMC samples).

