## Supplementary Text for "Negative Affect Induces Rapid Learning of Counterfactual Representations: A Model-based Facial Expression Analysis Approach"

Table S1

*All description-based games used in the current study*

| Set | Game | Option A | | | | Option B | | | | Pr(Choose A) | |
|---|---|---|---|---|---|---|---|---|---|---|---|
| | | $A_H$ | ($Pr_H$) | $A_L$ | ($Pr_L$) | $B_H$ | ($Pr_H$) | $B_L$ | ($Pr_L$) | No FB | FB |
| 1 | 1 | 3 | (1) | – | – | 4 | (.8) | 0 | (.2) | .41 | .16 |
| 1 | 2 | 3 | (.25) | 0 | (.75) | 4 | (.2) | 0 | (.8) | .25 | .13 |
| 1 | 3 | -1 | (1) | – | – | 0 | (.5) | -2 | (.5) | .28 | .1 |
| 1 | 4 | 1 | (1) | – | – | 2 | (.5) | 0 | (.5) | .5 | .18 |
| 1 | 5 | -3 | (1) | – | – | 0 | (.2) | -4 | (.8) | .31 | .18 |
| 1 | 6 | 0 | (.75) | -3 | (.25) | 0 | (.8) | -4 | (.2) | .46 | .31 |
| 1 | 7 | -1 | (1) | – | – | 0 | (.95) | -20 | (.05) | .38 | .12 |
| 1 | 8 | 1 | (1) | – | – | 20 | (.05) | 0 | (.95) | .47 | .34 |
| 1 | 9 | 1 | (1) | – | – | 100 | (.01) | 0 | (.99) | .38 | .34 |
| 1 | 10 | 2 | (1) | – | – | 101 | (.01) | 1 | (.99) | .32 | .33 |
| 1 | 11 | 19 | (1) | – | – | 20 | (.9) | -20 | (.1) | .78 | .53 |
| 1 | 12 | 0 | (1) | – | – | 50 | (.5) | -50 | (.5) | .55 | .22 |
| 1 | 13 | 0 | (1) | – | – | 50 | (.5) | -50 | (.5) | .5 | .22 |
| 1 | 14 | 0 | (1) | – | – | 1 | (.5) | -1 | (.5) | .38 | .29 |
| 1 | 15 | 7 | (1) | – | – | 50 | (.5) | 1 | (.5) | .09 | .01 |
| 1 | 16 | 7 | (1) | – | – | 50 | (.5) | -1 | (.5) | .16 | .02 |
| 1 | 17 | 30 | (1) | – | – | 50 | (.5) | 1 | (.5) | .63 | .32 |
| 1 | 18 | 30 | (1) | – | – | 50 | (.5) | -1 | (.5) | .65 | .3 |
| 1 | 24 | -2 | (1) | – | – | -1 | (.5) | -3 | (.5) | .38 | .19 |
| 1 | 25 | 2 | (1) | – | – | 3 | (.5) | 1 | (.5) | .49 | .26 |
| 1 | 26 | 16 | (1) | – | – | 50 | (.4) | 1 | (.6) | .3 | .06 |
| 1 | 28 | 6 | (.5) | 0 | (.5) | 9 | (.5) | 0 | (.5) | .02 | .01 |
| 1 | 29 | 2 | (1) | – | – | 3 | (1) | – | – | .02 | 0 |
| 1 | 30 | 6 | (.5) | 0 | (.5) | 8 | (.5) | 0 | (.5) | .02 | 0 |
| 2 | 32 | 24 | (.75) | -4 | (.25) | 82 | (.25) | 3 | (.75) | .16 | .06 |
| 2 | 33 | -3 | (1) | – | – | 14 | (.4) | -22 | (.6) | .51 | .28 |
| 2 | 35 | -5 | (1) | – | – | 47 | (.01) | -15 | (.99) | .69 | .73 |
| 2 | 37 | 23 | (.9) | 0 | (.1) | 64 | (.4) | -7 | (.6) | .38 | .21 |
| 2 | 38 | 24 | (1) | – | – | 34 | (.05) | 28 | (.95) | .05 | 0 |
| 2 | 43 | 14 | (1) | – | – | 12 | (.9) | 9 | (.1) | .8 | .83 |
| 2 | 44 | 23 | (1) | – | – | 24 | (.99) | -33 | (.01) | .6 | .31 |
| 2 | 46 | 37 | (.01) | 9 | (.99) | 30 | (.6) | -37 | (.4) | .65 | .31 |
| 2 | 51 | 42 | (.8) | -18 | (.2) | 68 | (.2) | 23 | (.8) | .1 | .04 |
| 2 | 52 | 46 | (.2) | 0 | (.8) | 46 | (.25) | -2 | (.75) | .51 | .41 |
| 2 | 53 | 28 | (1) | – | – | 42 | (.75) | -22 | (.25) | .49 | .26 |
| 2 | 54 | 18 | (1) | – | – | 64 | (.5) | -33 | (.5) | .47 | .31 |
| 2 | 56 | -8 | (1) | – | – | -5 | (.99) | -34 | (.01) | .14 | .02 |

| | | | | | | | | | | | |
|---|---|---|---|---|---|---|---|---|---|---|---|
| 2 | 58 | 85 | (.4) | -7 | (.6) | 40 | (.25) | 24 | (.75) | .23 | .17 |
| 2 | 59 | 17 | (.25) | 16 | (.75) | 43 | (.4) | 2 | (.6) | .25 | .12 |
| 2 | 60 | 51 | (.1) | 21 | (.9) | 38 | (.6) | 1 | (.4) | .38 | .32 |
| 3 | 62 | 25 | (1) | – | – | 45 | (.2) | 17 | (.8) | .48 | .28 |
| 3 | 65 | 12 | (.4) | -16 | (.6) | -5 | (1) | – | – | .49 | .2 |
| 3 | 67 | 85 | (.25) | 4 | (.75) | 54 | (.25) | 11 | (.75) | .41 | .2 |
| 3 | 68 | 12 | (1) | – | – | 102 | (.2) | -14 | (.8) | .49 | .19 |
| 3 | 70 | 18 | (1) | – | – | 35 | (.75) | -19 | (.25) | .46 | .11 |
| 3 | 71 | 13 | (.6) | -20 | (.4) | 76 | (.2) | -26 | (.8) | .44 | .21 |
| 3 | 72 | -9 | (1) | – | – | 13 | (.25) | -8 | (.75) | .11 | 0 |
| 3 | 75 | 13 | (1) | – | – | 50 | (.6) | -45 | (.4) | .46 | .19 |
| 3 | 77 | 1 | (1) | – | – | 38 | (.4) | -9 | (.6) | .21 | .02 |
| 3 | 78 | 19 | (1) | – | – | 44 | (.05) | 9 | (.95) | .8 | .62 |
| 3 | 79 | 32 | (.01) | 19 | (.99) | 65 | (.01) | 9 | (.99) | .7 | .79 |
| 3 | 80 | 3 | (1) | – | – | 50 | (.4) | -36 | (.6) | .42 | .16 |
| 3 | 83 | 9 | (1) | – | – | 64 | (.01) | 9 | (.99) | .08 | .01 |
| 3 | 84 | 27 | (1) | – | – | 22 | (.99) | -7 | (.01) | .85 | .92 |
| 3 | 85 | 20 | (1) | – | – | 70 | (.25) | 6 | (.75) | .34 | .14 |
| 3 | 89 | 17 | (1) | – | – | 44 | (.1) | 17 | (.9) | .09 | 0 |
| 3 | 90 | 10 | (1) | – | – | 31 | (.75) | -49 | (.25) | .41 | .1 |
| 4 | 91 | 7 | (1) | – | – | 16 | (.1) | 10 | (.9) | .04 | 0 |
| 4 | 92 | 8 | (.8) | -37 | (.2) | 102 | (.2) | -29 | (.8) | .36 | .18 |
| 4 | 94 | 7 | (1) | – | – | 6 | (.75) | 1 | (.25) | .78 | .89 |
| 4 | 96 | 35 | (.5) | -47 | (.5) | -10 | (.75) | -15 | (.25) | .45 | .24 |
| 4 | 97 | 10 | (1) | – | – | 45 | (.2) | -5 | (.8) | .59 | .32 |
| 4 | 100 | 18 | (.6) | -29 | (.4) | -1 | (1) | – | – | .22 | .1 |
| 4 | 104 | -6 | (1) | – | – | 3 | (.99) | -27 | (.01) | .05 | .01 |
| 4 | 105 | 30 | (1) | – | – | 90 | (.01) | 36 | (.99) | .06 | 0 |
| 4 | 108 | 16 | (1) | – | – | 91 | (.2) | -11 | (.8) | .52 | .34 |
| 4 | 109 | 11 | (1) | – | – | 26 | (.5) | -9 | (.5) | .41 | .2 |
| 4 | 111 | 28 | (1) | – | – | 47 | (.6) | -13 | (.4) | .52 | .16 |
| 4 | 114 | 72 | (.01) | -2 | (.99) | 112 | (.25) | -33 | (.75) | .31 | .14 |
| 4 | 117 | -6 | (1) | – | – | 7 | (.5) | -30 | (.5) | .48 | .21 |
| 4 | 120 | -9 | (.95) | -26 | (.05) | -1 | (.1) | -11 | (.9) | .29 | .2 |
| 5 | 122 | 68 | (.05) | -14 | (.95) | -11 | (.9) | -36 | (.1) | .46 | .28 |
| 5 | 123 | 28 | (.75) | -13 | (.25) | 57 | (.1) | 16 | (.9) | .08 | .02 |
| 5 | 124 | 15 | (.95) | 7 | (.05) | 42 | (.01) | 19 | (.99) | .06 | 0 |
| 5 | 125 | 28 | (1) | – | – | 41 | (.4) | 12 | (.6) | .44 | .22 |
| 5 | 128 | -3 | (1) | – | – | 32 | (.4) | -16 | (.6) | .15 | .04 |
| 5 | 130 | 72 | (.4) | -41 | (.6) | 16 | (.01) | 1 | (.99) | .21 | .1 |

| | | | | | | | | | | | |
|---|---|---|---|---|---|---|---|---|---|---|---|
| 5 | 131 | 18 | (1) | – | – | 45 | (.01) | 11 | (.99) | .65 | .65 |
| 5 | 135 | 6 | (1) | – | – | 8 | (.5) | -1 | (.5) | .55 | .35 |
| 5 | 136 | 4 | (1) | – | – | 25 | (.01) | -5 | (.99) | .72 | .75 |
| 5 | 138 | 23 | (1) | – | – | 21 | (.8) | 16 | (.2) | .78 | .8 |
| 5 | 140 | -2 | (1) | – | – | 9 | (.25) | 8 | (.75) | .01 | 0 |
| 5 | 141 | 28 | (.8) | -26 | (.2) | 22 | (.75) | 2 | (.25) | .09 | .01 |
| 5 | 142 | 23 | (1) | – | – | 29 | (.8) | -8 | (.2) | .48 | .15 |
| 5 | 143 | 67 | (.5) | -39 | (.5) | 93 | (.25) | -15 | (.75) | .25 | .02 |
| 6 | 157 | 16 | (1) | – | – | 33 | (.5) | 14 | (.5) | .03 | 0 |
| 6 | 159 | 14 | (1) | – | – | 20 | (.95) | 16 | (.05) | .03 | 0 |
| 6 | 160 | 60 | (.1) | 19 | (.9) | 34 | (.8) | 23 | (.2) | .06 | .01 |
| 6 | 162 | 39 | (.6) | 14 | (.4) | 48 | (.25) | 24 | (.75) | .19 | .07 |
| 6 | 163 | 25 | (1) | – | – | 35 | (.2) | 12 | (.8) | .75 | .49 |
| 6 | 164 | 1 | (.5) | 1 | (.5) | 40 | (.01) | -4 | (.99) | .77 | .61 |
| 6 | 166 | -6 | (1) | – | – | 15 | (.6) | -10 | (.4) | 0 | 0 |
| 6 | 167 | 13 | (1) | – | – | 12 | (.8) | 8 | (.2) | .86 | .9 |
| 6 | 172 | 21 | (1) | – | – | 26 | (.8) | -24 | (.2) | .63 | .28 |
| 7 | 182 | 28 | (1) | – | – | 27 | (.95) | -11 | (.05) | .9 | .94 |
| 7 | 185 | 7 | (1) | – | – | 18 | (.99) | -19 | (.01) | .08 | 0 |
| 7 | 186 | 19 | (1) | – | – | 97 | (.1) | 5 | (.9) | .5 | .23 |
| 7 | 187 | -3 | (1) | – | – | 43 | (.2) | -20 | (.8) | .44 | .21 |
| 7 | 189 | 2 | (1) | – | – | 4 | (.75) | -23 | (.25) | .65 | .34 |
| 7 | 190 | 10 | (1) | – | – | 28 | (.2) | -1 | (.8) | .62 | .28 |
| 7 | 191 | 16 | (1) | – | – | 15 | (.8) | -12 | (.2) | .89 | .91 |
| 7 | 192 | 10 | (1) | – | – | 59 | (.01) | 4 | (.99) | .68 | .49 |
| 7 | 200 | 16 | (1) | – | – | 40 | (.6) | -15 | (.4) | .39 | .09 |
| 7 | 202 | 53 | (.1) | 27 | (.9) | 32 | (.95) | 9 | (.05) | .25 | .15 |
| 7 | 207 | 22 | (.9) | -41 | (.1) | 14 | (.99) | 6 | (.01) | .19 | .12 |

*Note*. Each of the description-based games used in study 1 are shown here, along with the set of games that they belong to. $A_H$ = high outcome for option A. $A_L$ = low outcome for option A (same notation for option B). $Pr_H$ = high outcome for given option. $Pr_L$ = low outcome for given option. Pr(Choose A) indicates the portion of trials that subjects choose option A both in the 5 trials before feedback (No FB) and in the remaining 20 trials following feedback (FB).

Figure S1.

*Posterior predictive simulations for set 1, games 1-6*

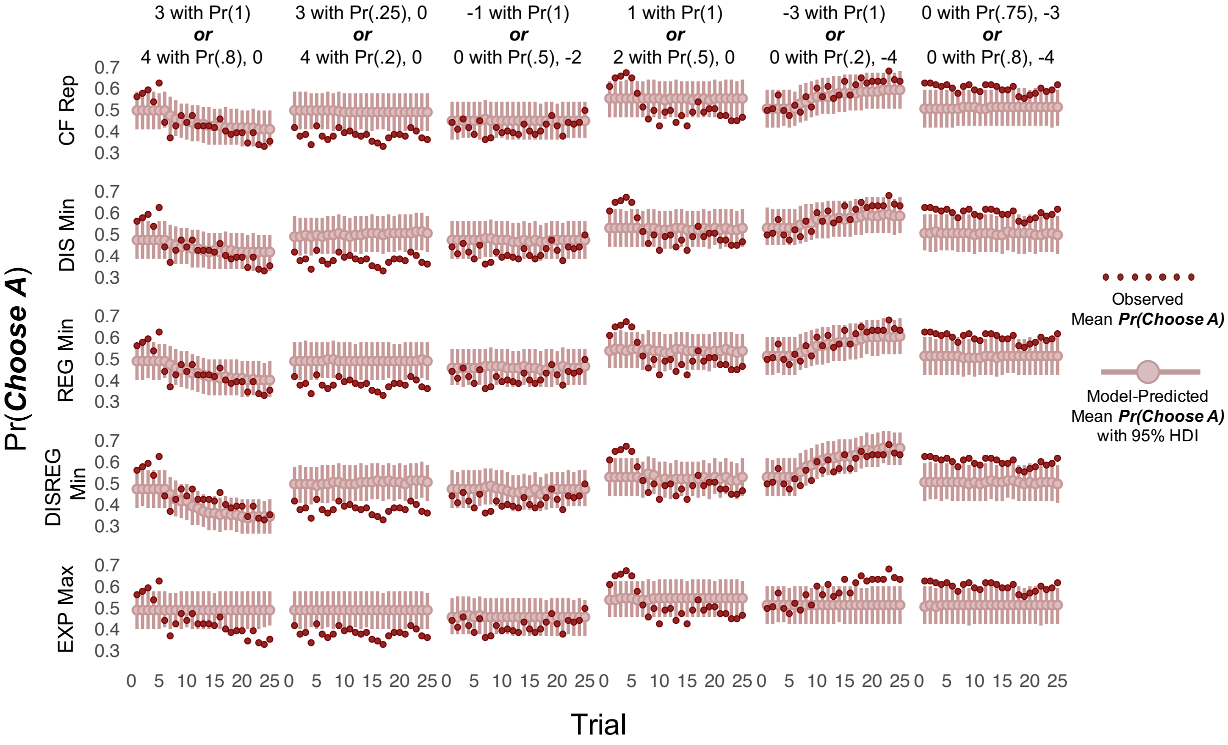


Figure S2.

*Posterior predictive simulations for set 1, games 7-12*

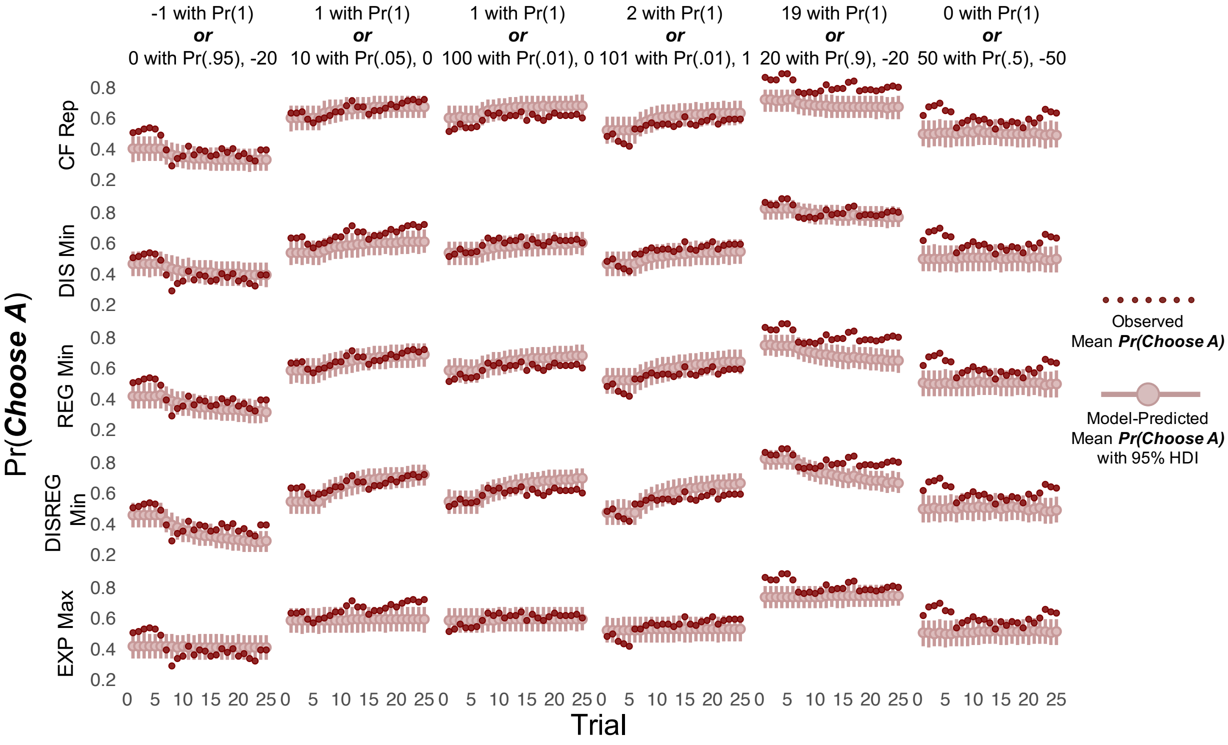

Figure S3.

*Posterior predictive simulations for set 1, games 13-18*

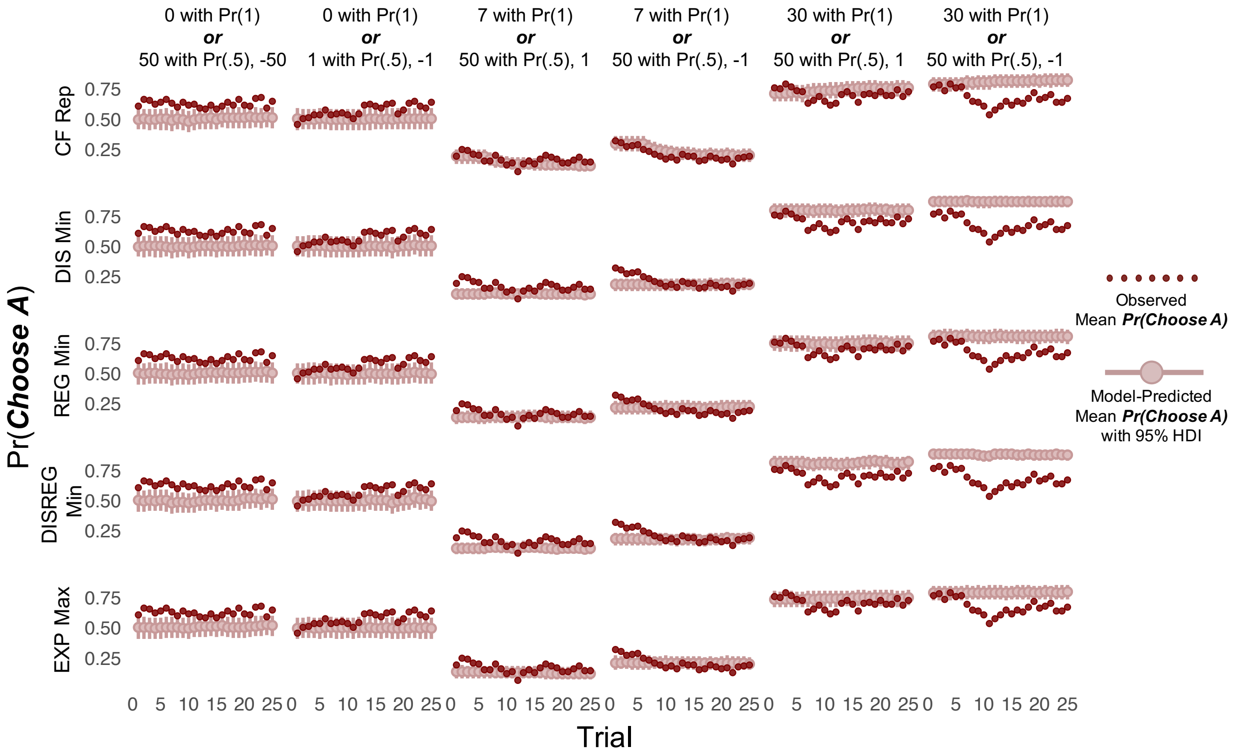


Figure S4.

*Posterior predictive simulations for set 1, games 24-26 and 28-30*

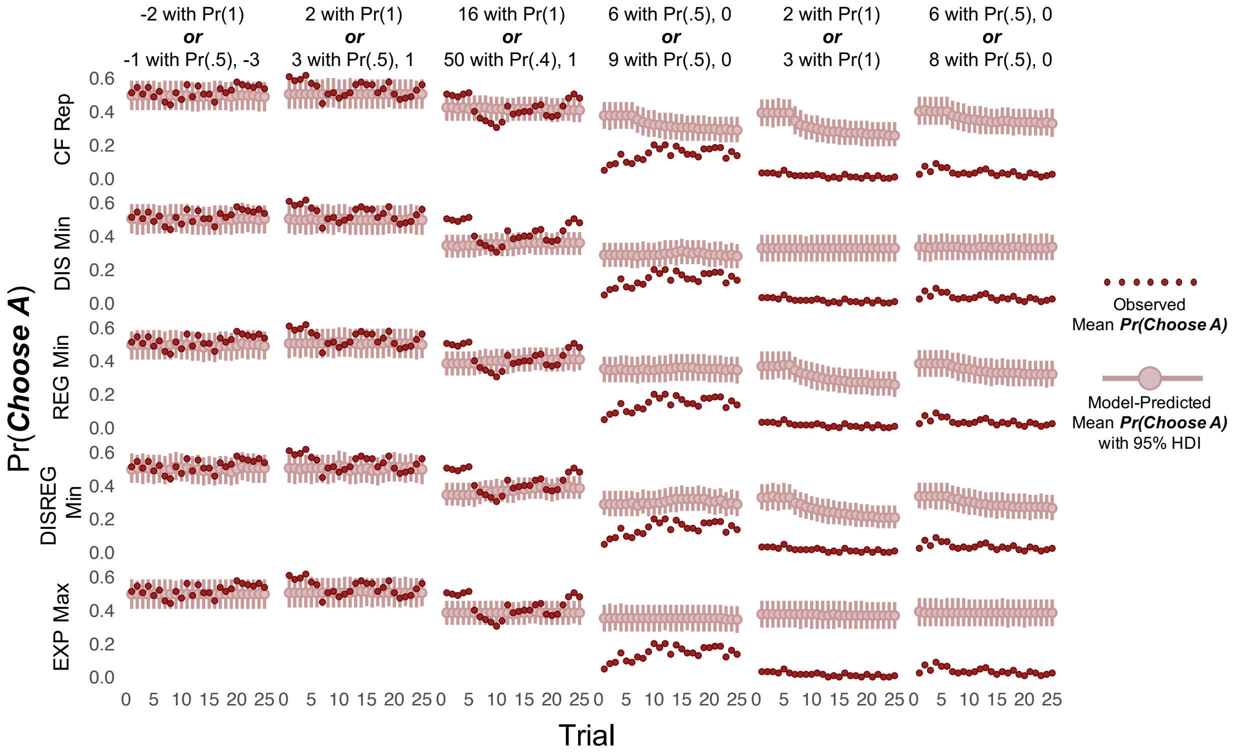

Figure S5.

*Posterior predictive simulations for set 2, games 32, 33, 35, 37, 38, and 43*

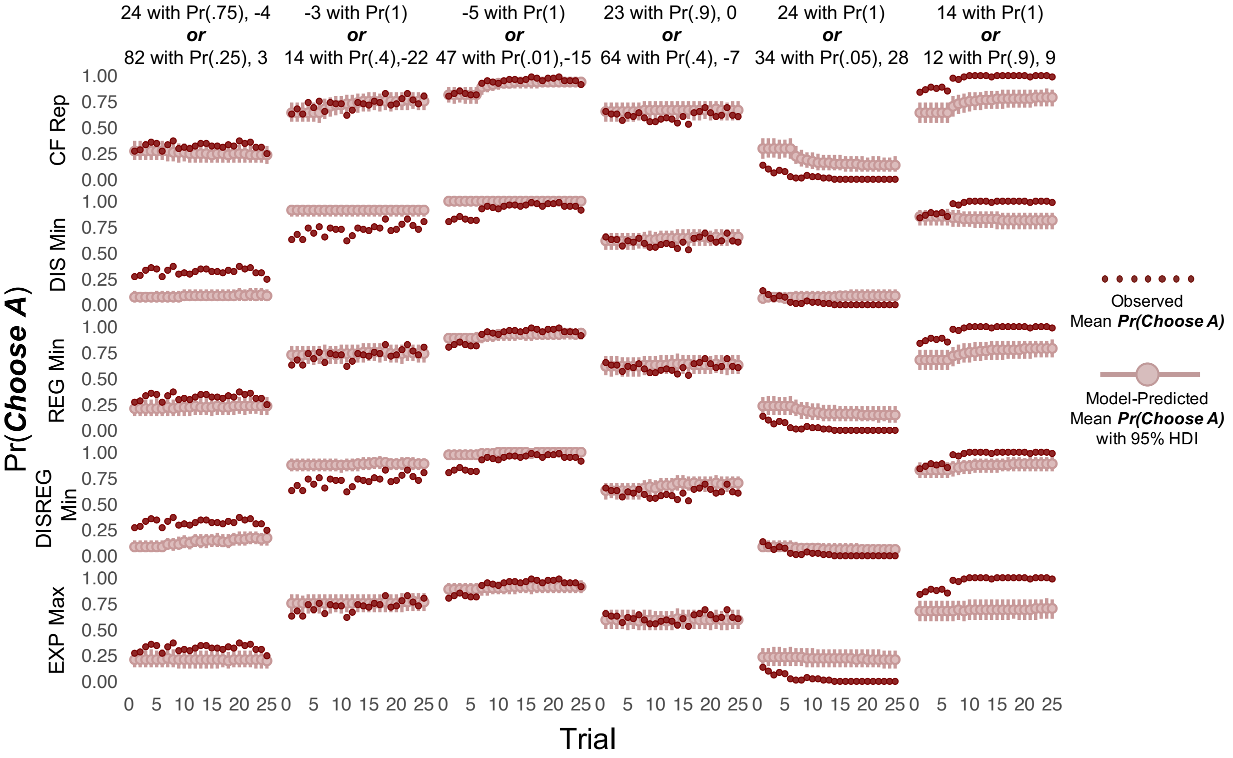


Figure S6.

*Posterior predictive simulations for set 2, games 44, 46, 51, and 52-54*

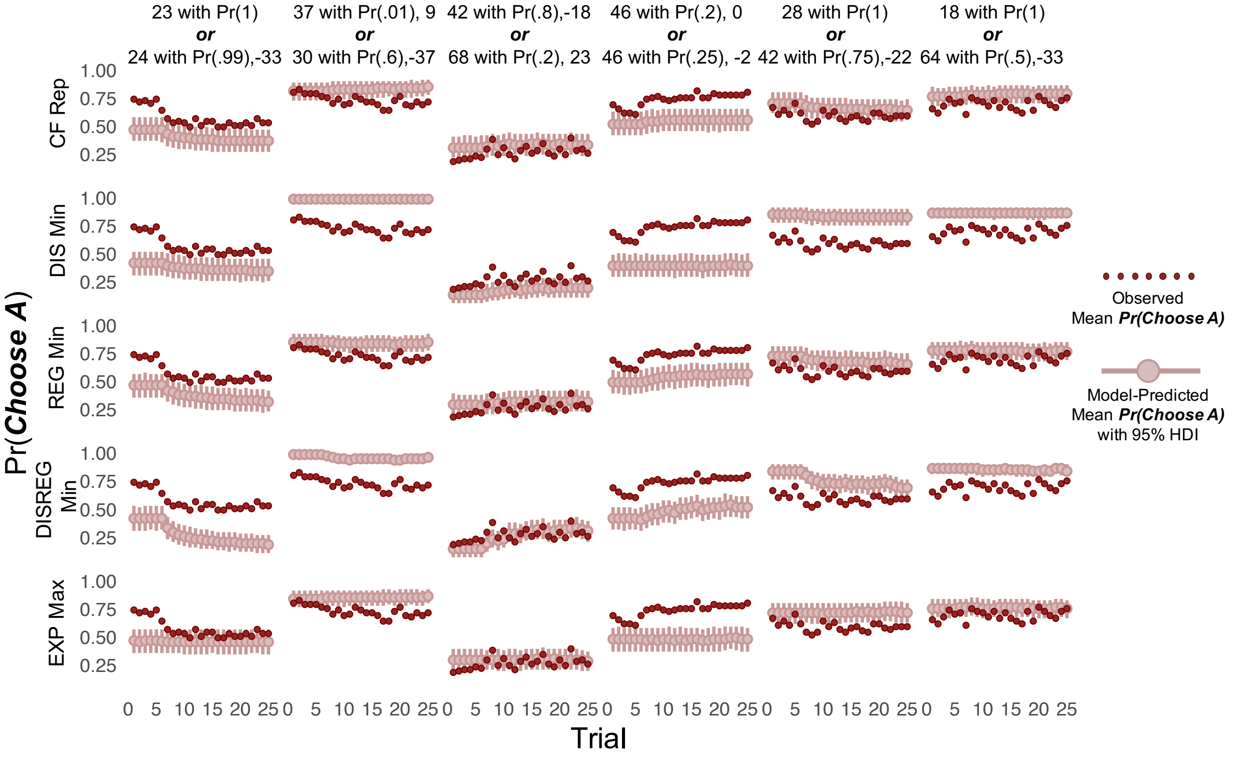

Figure S7.

*Posterior predictive simulations for set 2, games 56, and 58-60*

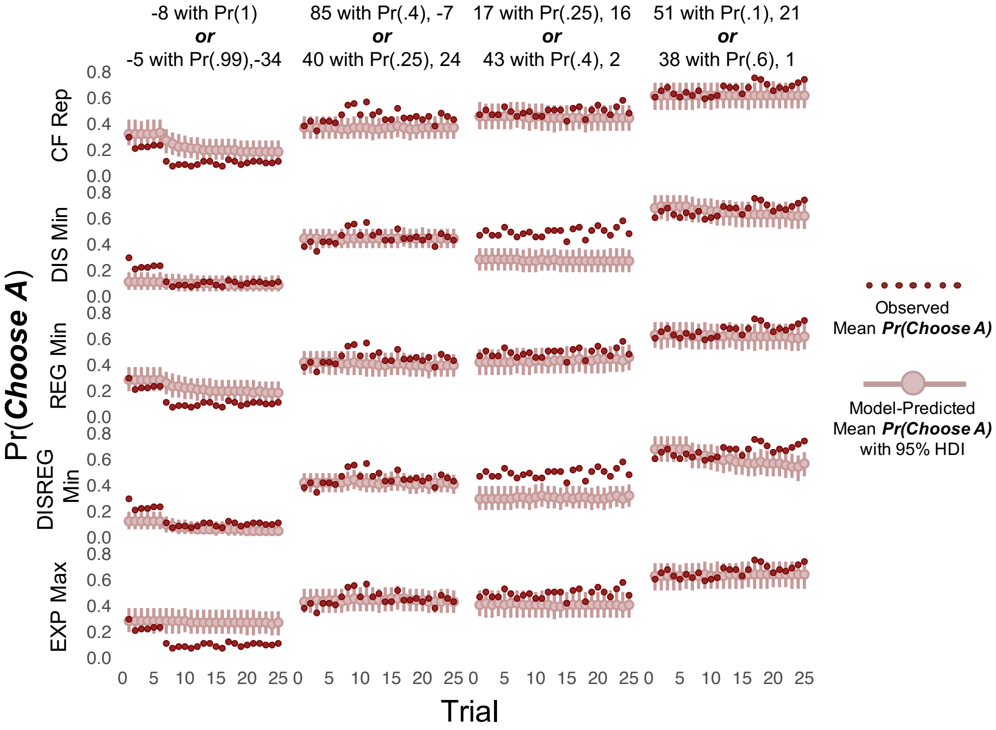


Figure S8.

*Posterior predictive simulations for set 3, games 62, 65, 67, 68, 70, and 71*

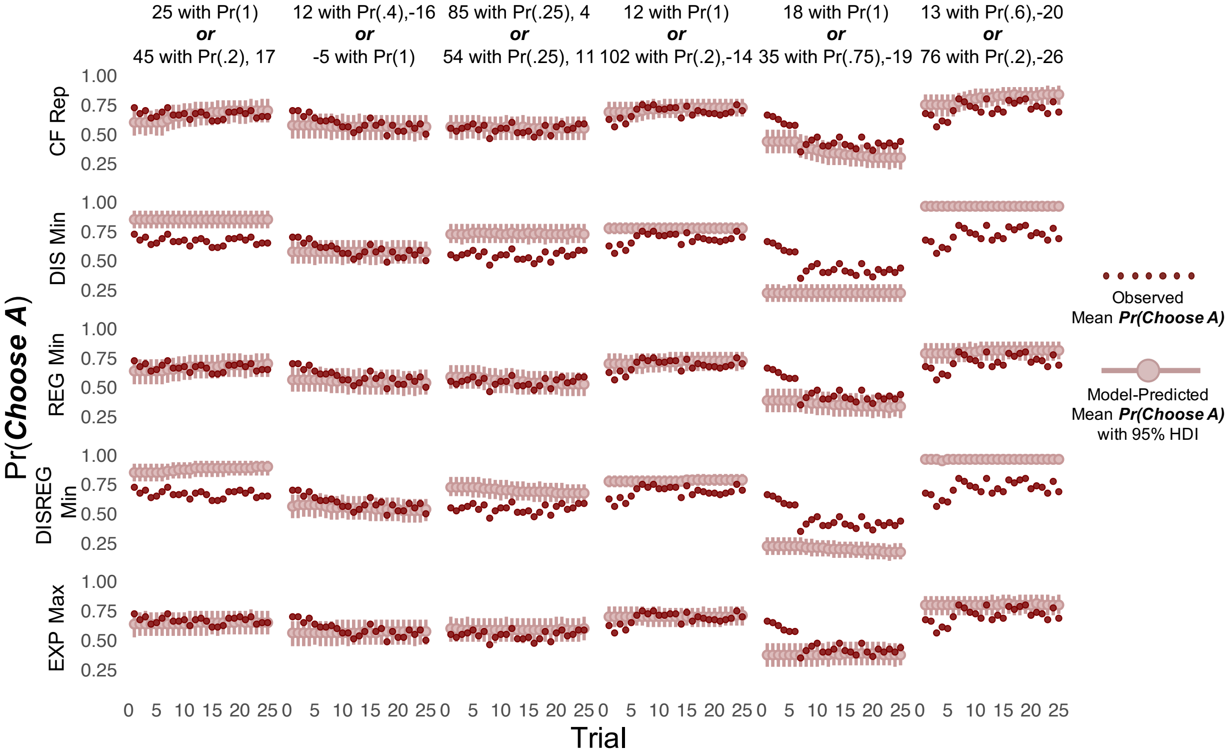

Figure S9.

*Posterior predictive simulations for set 3, games 72, 75, 77, and 78-80*

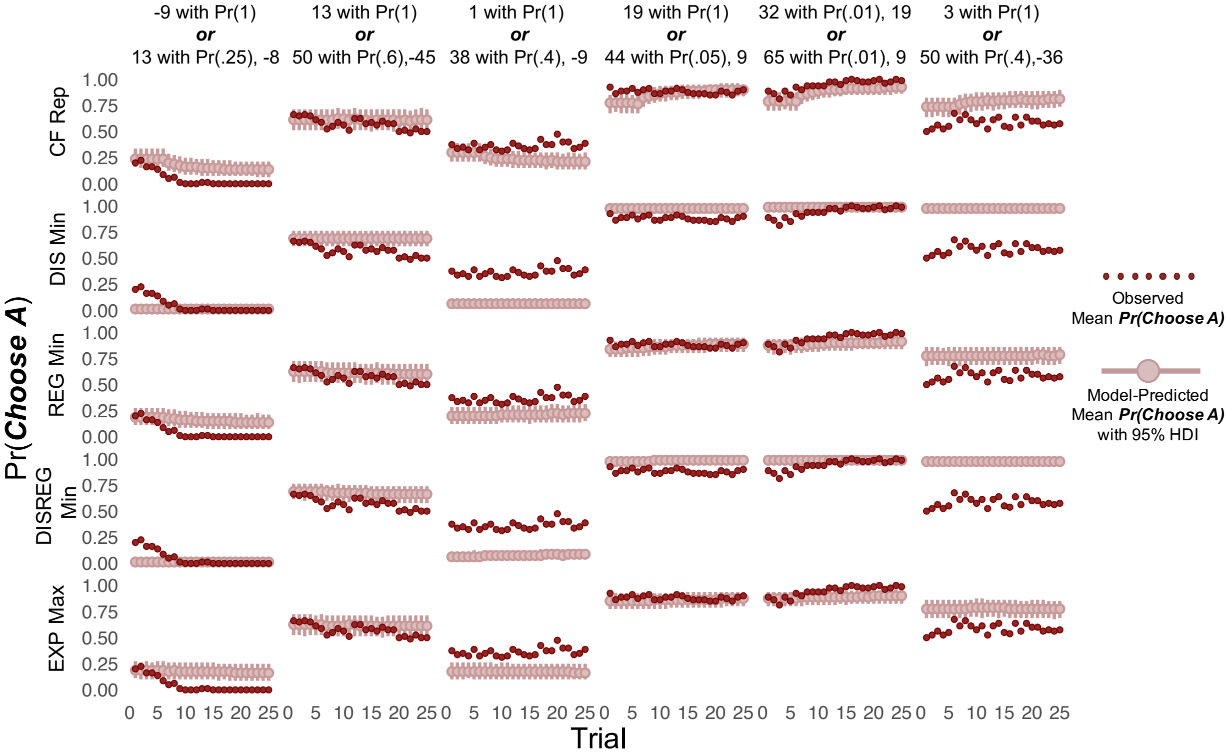


Figure S10.

*Posterior predictive simulations for set 3, games 83-85, 89, and 90*

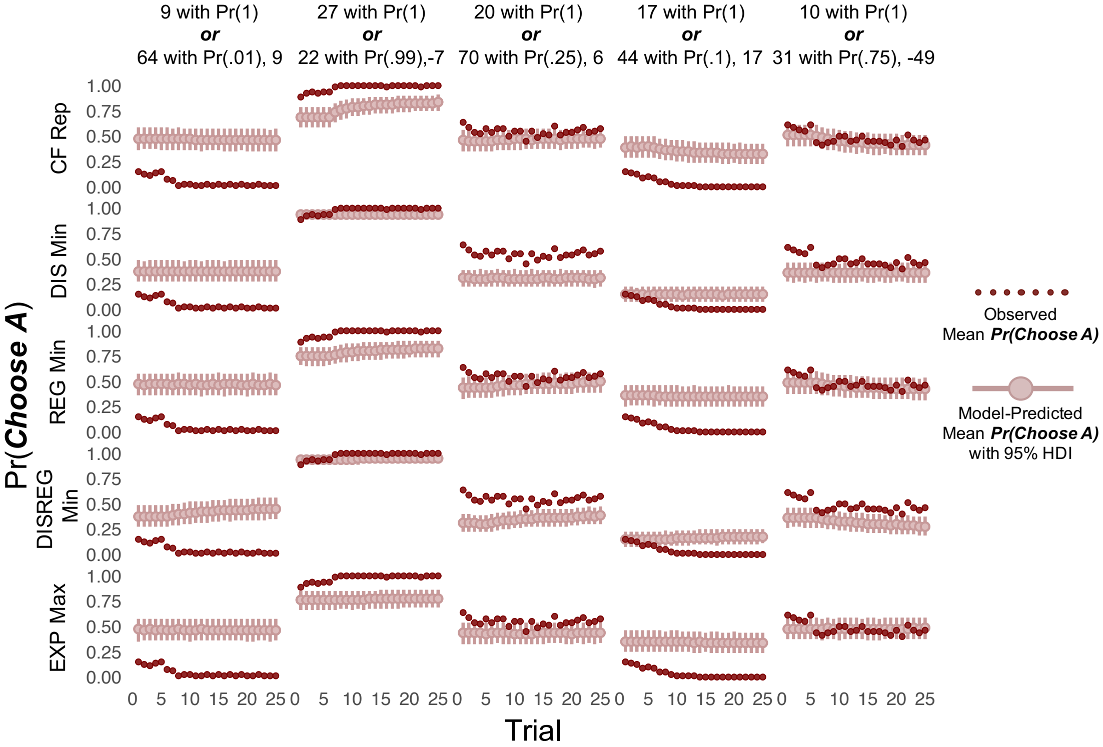

Figure S11.

*Posterior predictive simulations for set 4, games 91, 92, 94, 96, 97, and 100*

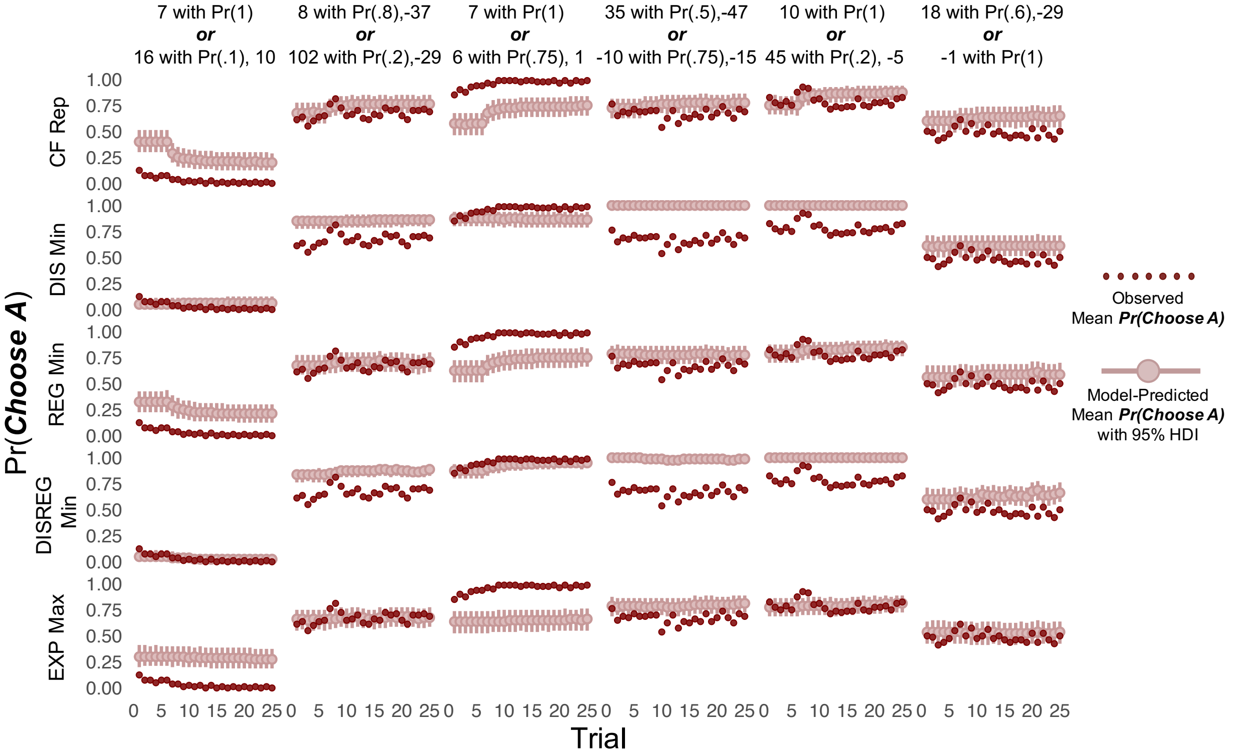


Figure S12.

*Posterior predictive simulations for set 4, games 104, 105, 108, 109, 111, and 114*

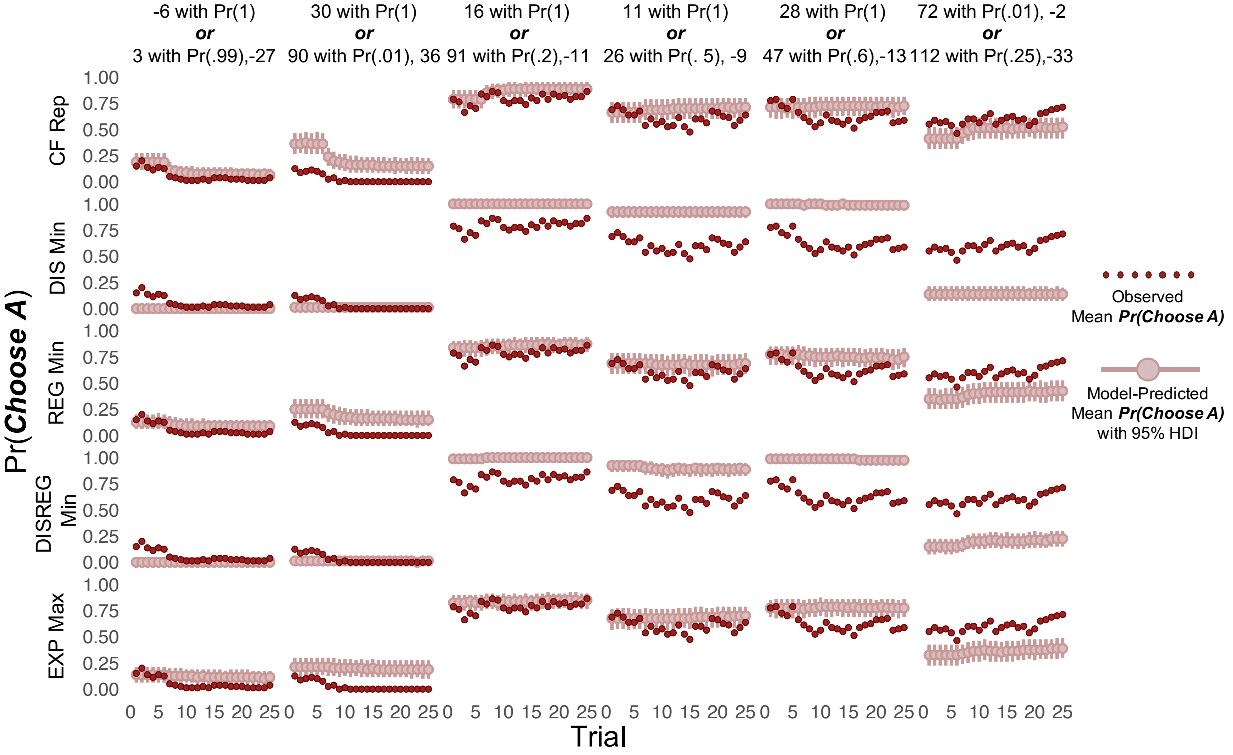

Figure S13.

*Posterior predictive simulations for set 4, games 117 and 120*

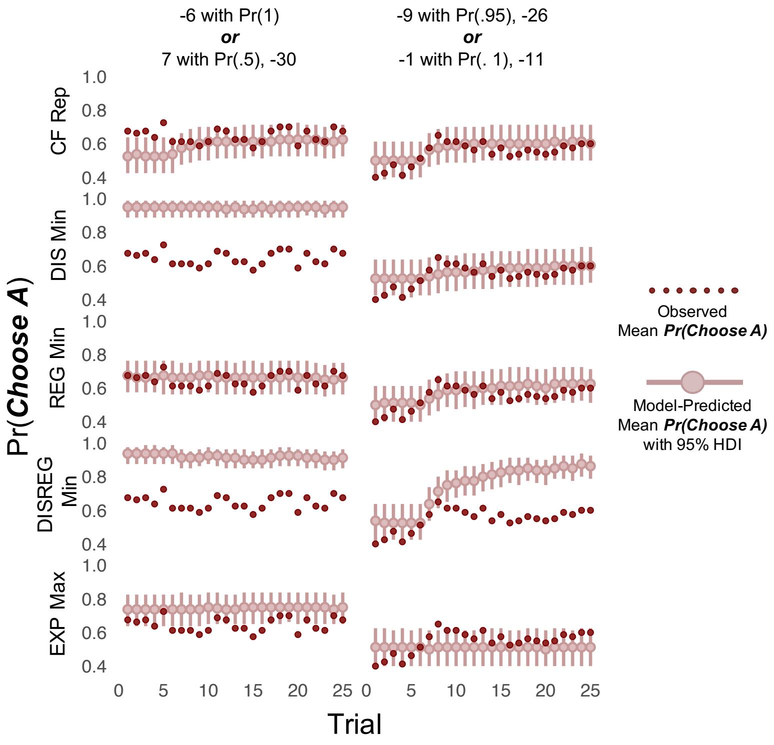


Figure S14.

*Posterior predictive simulations for set 4, games 122-125, 128, and 130*

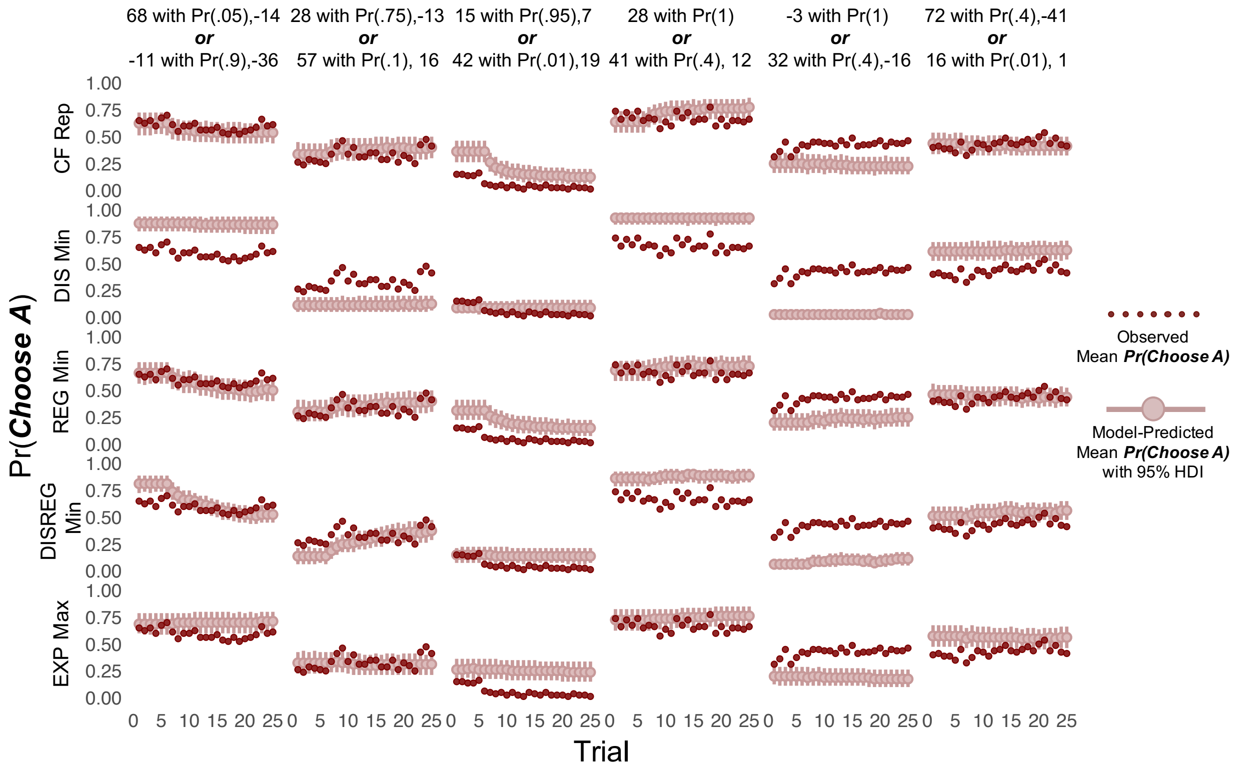

Figure S15.

*Posterior predictive simulations for set 5, games 131, 135, 136, 138, 140, and 141*

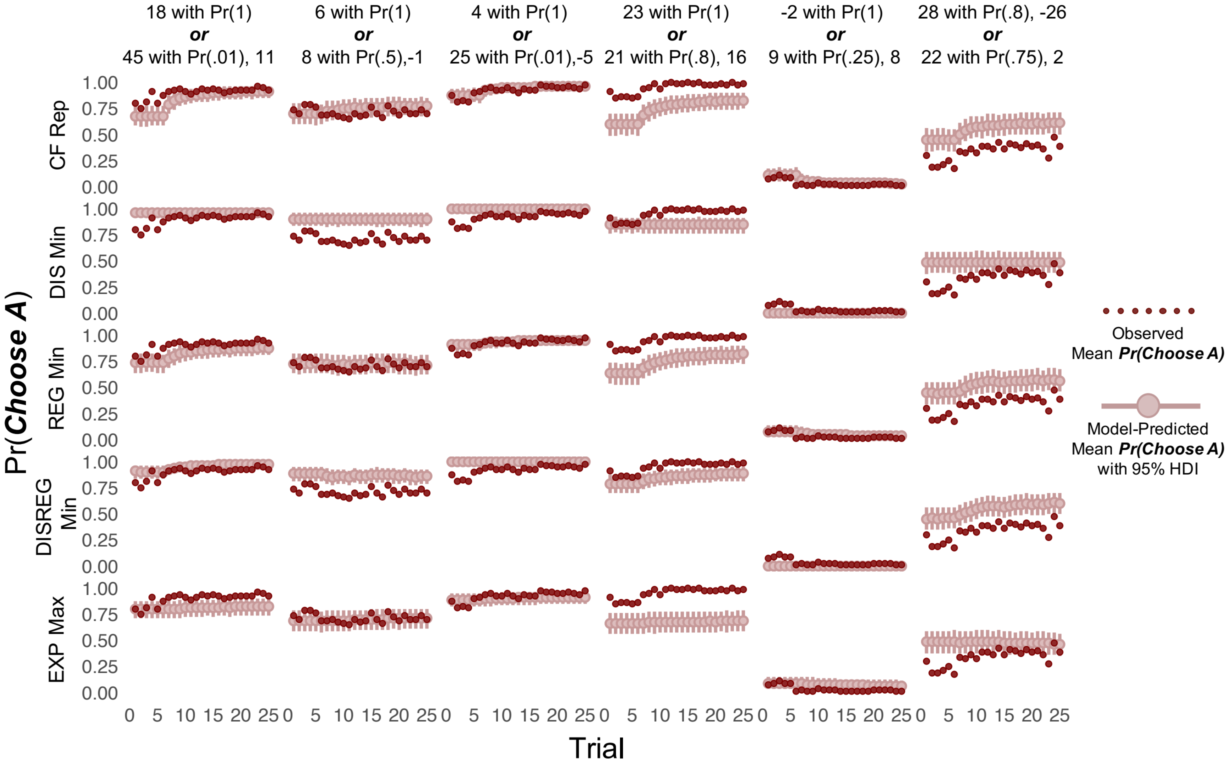


Figure S16.

*Posterior predictive simulations for set 5, games 142 and 143*

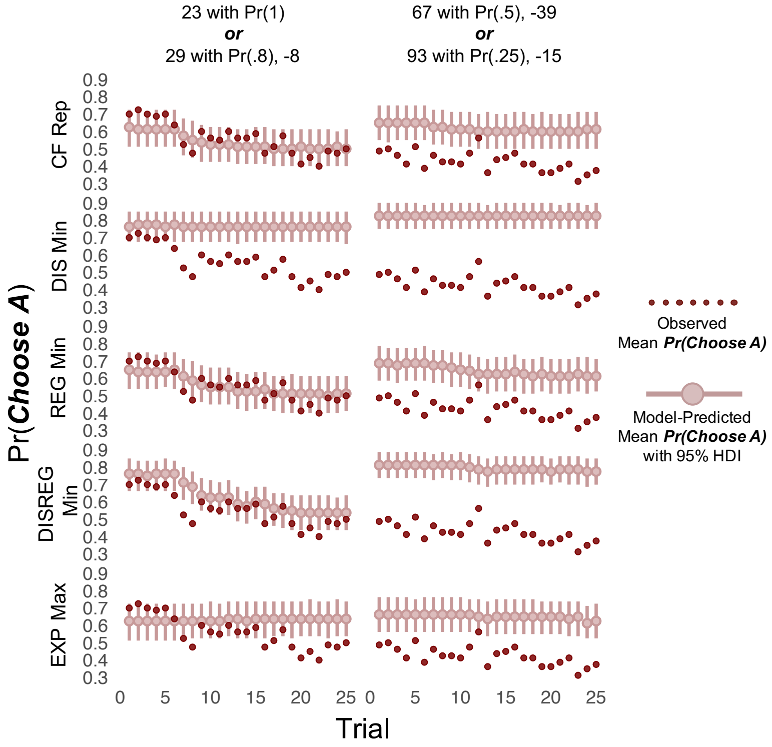

Figure S17.

*Posterior predictive simulations for set 6, games 157, 159, 160, 162, 163, and 164*

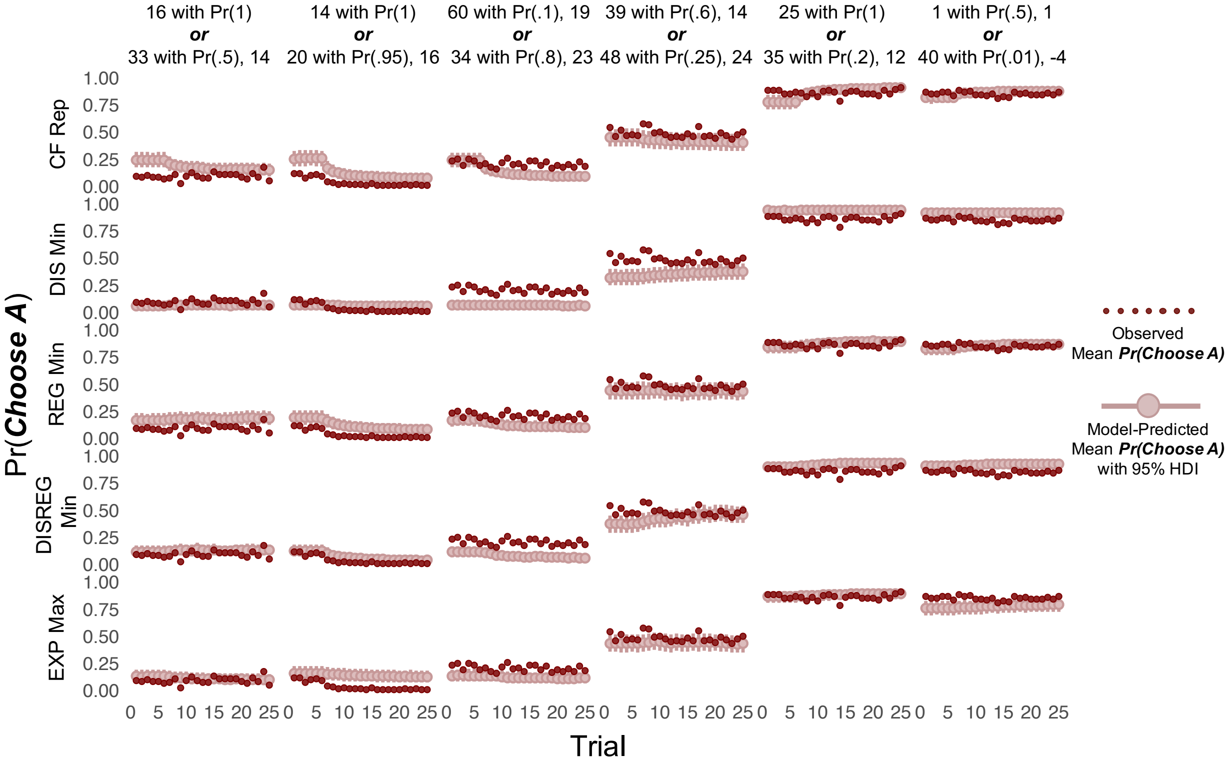


Figure S18.

*Posterior predictive simulations for set 6, games 166, 167, and 172*

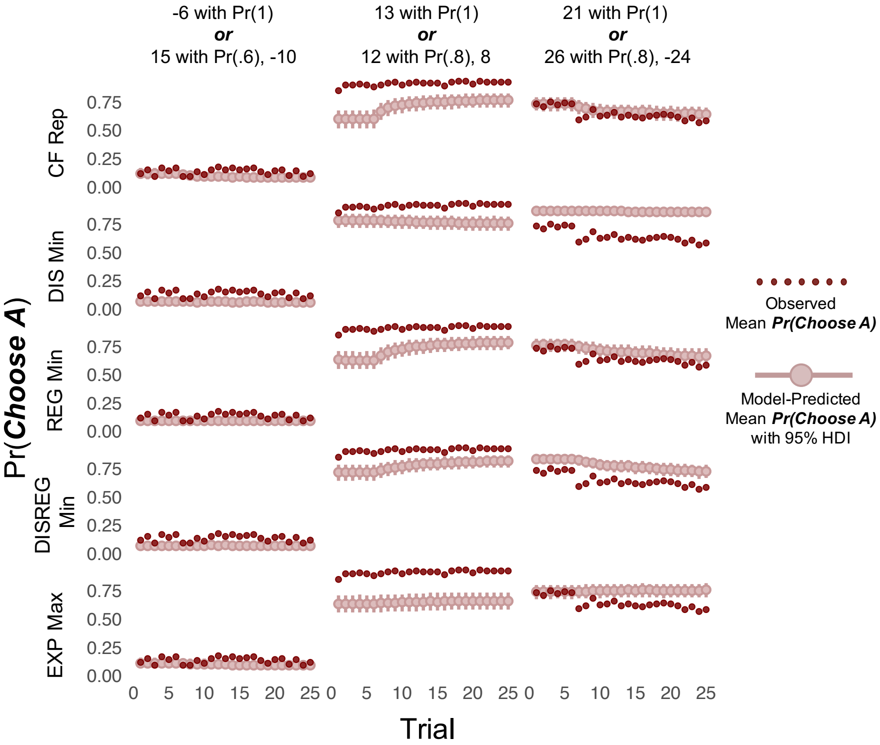

Figure S19.

*Posterior predictive simulations for set 7, games 182, 185-187, 189, and 190*

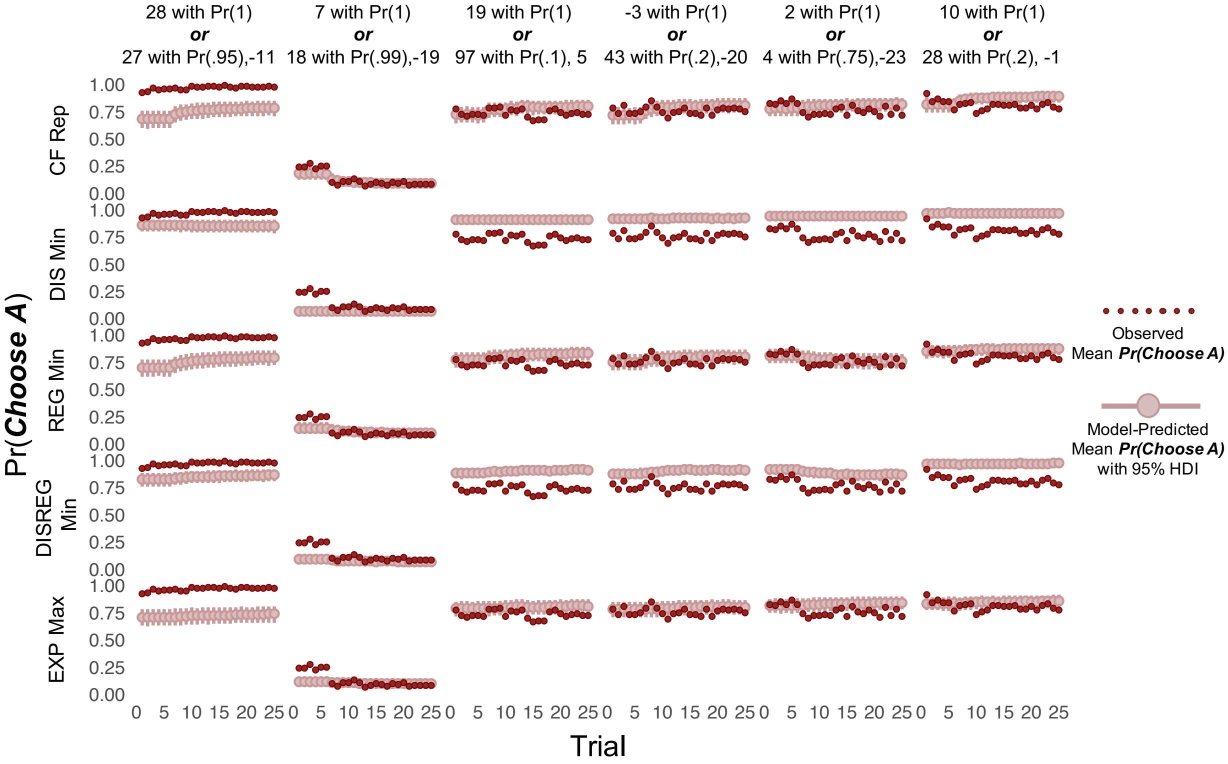


Figure S20.

*Posterior predictive simulations for set 7, games 191, 192, 200, 202, and 207*

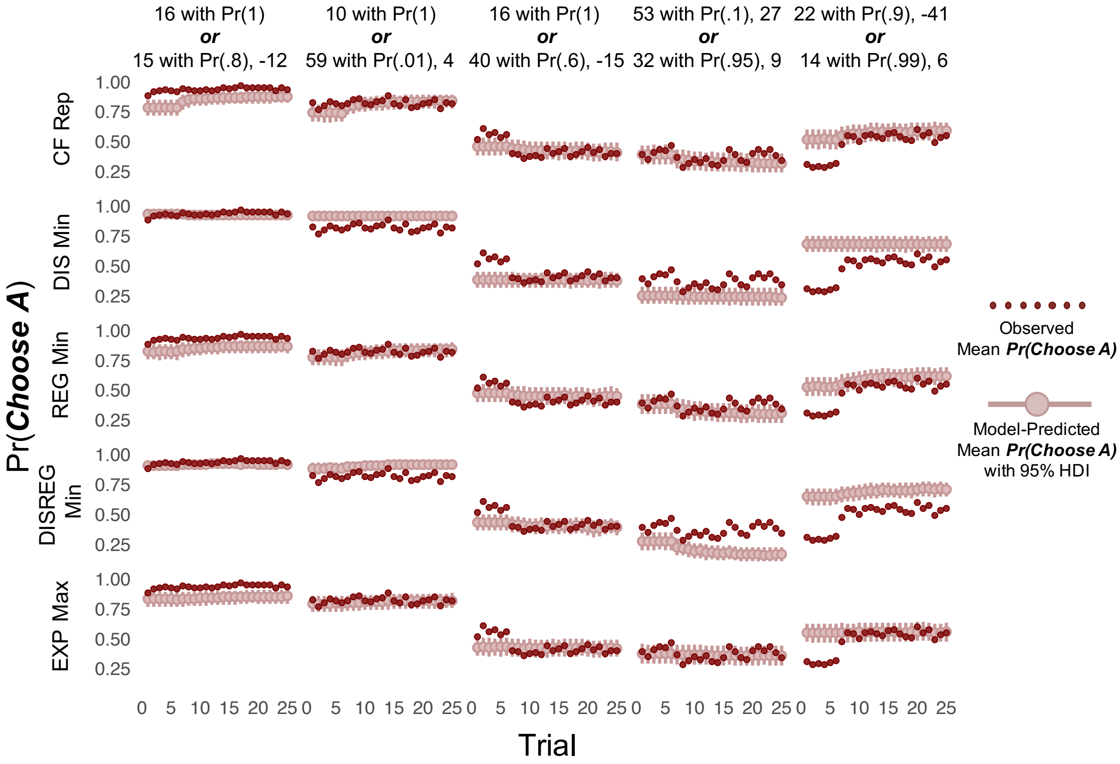
